# Alterations of the composition and spatial organization of the microenvironment following non-dysplastic Barrett’s esophagus through progression to cancer

**DOI:** 10.1101/2025.06.09.657642

**Authors:** Meng-Lay Lin, John W. Hickey, Adam Passman, Emanuela Carlotti, Shruthi Devkumar, Yannick H Derwa, Joanne ChinAleong, Richard J. Hackett, James Evans, Veena Sangwan, Marco R. Novelli, Manuel Rodriguez Justo, Nicholas A. Wright, Marnix Jansen, Tim J Underwood, Sui Huang, Philippe Gascard, Mairi H. McLean, Christian M. Schürch, Thea D. Tlsty, Lorenzo Ferri, Garry P. Nolan, Stuart A.C. McDonald

## Abstract

Barrett’s esophagus (BE), a metaplastic condition that is the only known precursor for esophageal adenocarcinoma (EAC), is relatively common, but progression to cancer is infrequent. BE is inflamed but the contribution of the immune system to the carcinogenic process is unknown. To this end, we contrasted non-dysplastic metaplasia of BE patients, captured when they did not progress (non-progressors), did subsequently, but had not yet progressed (pre-progressors) or had already progressed to EAC (progressors). Using spatial multiplexed 56-protein analysis, serial laser capture microdissection (LCM) RNAseq and shallow whole genome sequencing, we identified prooncogenic immune neighbourhoods and dysregulated immune cell populations predictive of subsequent progression to EAC. Indeed, spatial analysis revealed that M1 macrophages, regulatory natural killer (NK) cells, neutrophils and altered ratios of intraepithelial CD4^+^ and CD8^+^ lymphocytes typify tumor microenvironmental (TME) changes associated with cancer initiation. Spatially derived cell-to-cell interactions revealed progression-specific immune cell interaction signatures predominantly involving M1 macrophages NK cells and plasma cells. Furthermore, LCM RNAseq analysis identified gene expression ‘hot’ signatures enriched in pre-progression and progression samples. Notably, we also observed a correlation between immune cells and copy number alterations in progressor metaplasia. By exposing coordinated changes in the immune cell landscape in patients at high risk of developing EAC, this multi-omic dataset provides novel diagnostic and therapeutic opportunities

## Introduction

Metaplasia, induced by chronic inflammation, is a common adaptive phenotypic cell conversion which is considered as a facultative precancerous condition ^1^. Despite the known role of chronic inflammation in tumorigenesis^2^, how the interplay of its cellular components, notably the interactions between stromal and immune cells, drive progression of metaplastic epithelial cells to cancer, remains elusive. In this case, chronic inflammation may not reflect a failure of immune cells to repair damaged tissue but rather their inability to resolve inflammation, leading to persistent remodeling. This sustained dysregulation causes immunosuppression, resulting in immune escape of altered cells, as well as cellular stress responses which predispose to neoplastic transformation^3^.

Barrett’s esophagus (BE) is an example of a metaplastic state and is the only known precursor to esophageal adenocarcinoma (EAC)^4^. Ulceration due to exposure of the esophageal squamous epithelium to acid and bile reflux triggers a repair response from the adjacent gastric epithelium, which promotes metaplasia^1,5^. The genetic hallmarks of BE progression have been extensively studied^6–10^, demonstrating that progressive, yet non-dysplastic BE contains a higher diversity of copy number variations (CNVs). However, despite high mutation rates, no consistent driver mutations, aside from those affecting TP53^11–14^, have been reported in Barrett’s.

Relatively little is known about how the immune response and stroma change during the progression of metaplastic BE cells to cancer ^15,16^. Non-progressive BE has been associated with a Th2-dominant immune response^17^ which appears to shift to a Th1-like profile as BE progresses to EAC ^18^. Moreover, an increase in FOXP3^+^ CD4^+^ T cells and CD163^+^ myelomonocytic cells has been observed in BE progression^19,20^. Other studies, all with a focus on the transition from BE to dysplasia and cancer, have shown a loss of CD8^+^ T cells^20^. Our own previous scRNAseq studies are consistent with these published studies and, additionally, have shown that metaplasia is also marked by the appearance of fibroblasts with gastric identities and characteristics reminiscent of carcinoma-associated fibroblasts (CAFs)^21^. What is unclear is whether these changes in the metaplastic environment can already be detected in those patients prior to progression to dysplasia and cancer, key information which could then be used as a prognostic biomarker in surveillance.

Our primary aim in this work was to investigate how the stromal landscape in BE metaplastic lesions change before and at the time of cancer development. Our findings identify specific signatures for BE patients that are at risk of progressing to EAC. Defining microenvironments that promote progression of BE to cancer is critical^5^. To date, studies on prospective progressor (pre-progressor, where patient samples were taken years prior to cancer development) non-dysplastic BE (NDBE) have primarily focused on the genomic landscape which has revealed altered genotypes in patients, years before progression^7,8,10,22^. Here, we take advantage of a unique sample set afforded by surveillance program NDBE of demonstrated non-progressor, pre-progressor and proven progressor (non-dysplastic metaplasia from patients with diagnosed cancer or dysplasia) patients, to investigate combining transcriptomic, proteomic and genomic approaches, a strategy that allow us, for the first time, to ascertain the actual contribution and timeline of alterations of the stromal landscape associated with progression of NDBE to cancer.

Our data shows that an increase in immunoregulatory NK cells, neutrophils and M1 macrophages along with their spatial relationships with other immune cells such as Tregs and plasma cells represent an important immunological event in the progression to EAC. Additionally, altered ratios of CD4 to CD8 T cells in the BE epithelium is dependent on whether a BE patient will progress. Our work suggests that early changes in this tissue at the metaplastic stage may stratify risk of progression for BE patients, discriminating those patients likely to remain non-dysplastic from those that will progress. The early reprogramming of the immune and stromal compartments in metaplasia likely opens new avenues for risk stratification tools and potential therapeutic targets for preventing malignancy.

## Methods

### Patient Biopsies

Formalin-fixed paraffin-embedded (FFPE) archival biopsies were collected from three cohorts of BE patients for this study: (i) NP: Non-progressors (n=48), defined as patients who have no history of previous dysplasia or cancer for 7 years prior to the date of the study and have undergone at least 3 endoscopies; (ii) PP: Pre-progressors (n=26), defined as patients who have undergone at least two dysplasia/cancer-free endoscopies over a period of at least 7 years, the latest endoscopy being performed at least 2 years prior to the development of dysplasia or esophageal adenocarcinoma (EAC); (iii) P: Progressors (n=27), defined as patients who have developed EAC but whose histology revealed adjacent areas of NDBE. Anonymized metadata for all experiments are described in Supplemental data Spreadsheet S1. Only a portion of patient samples were used for each technique, and these are detailed later. Patient biopsies were sourced from tissue banks and hospital pathology archives (Barts Health Foundation Trust, University College Hospitals London Foundation Trust, Grampian NHS Trust) under ethical approval from Stanmore Research Ethics Committee 11/LO/1613 or East London Research Ethics committee 15/LO/1217. Patient biopsies were also collected from McGill University Health Centre (Montreal, Canada) under approval REB# 2007-856. Histological diagnoses were confirmed by at least 3 of 6 expert pathologists (JCA, MRN, MRJ, MJ, NAW and CMS) in a blinded fashion.

### Tissue microarrays (TMA) and virtual cores

Tissue microarrays were constructed using a maximized number of 0.6 mm cores targeted from a reference H&E section and evenly distributed across approximately 4 FFPE blocks per sample. For the PP cohort, only tissue sections were available for each patient and TMAs could not be constructed. Instead, a maximal number of “virtual” cores were taken on the entire section using a custom interactive web application developed with Shiny^9^, incorporating the Plotly^10^, ggforce^11^, and ggplot2^12^ R packages. To enable direct comparison with tissue microarray (TMA) cores, which have a diameter of 0.6 mm, the virtual core radius was set to 0.6 mm, based on a pixel-to-millimeter conversion rate of 0.5 µm/pixel. This approach ensured consistency in core size and scale, facilitating direct histological comparison between the virtual tissue section cores and traditional TMA cores. Virtual cores were representative of the metaplastic regions of physical cores as defined by the presence of NDBE. Overall, cell content of virtual cores between non-progressors and pre-progressors at the metaplastic stage was not found to be significantly different from physical cores (data not shown), thus ensuring the legitimacy of comparing virtual and physical cores.

### CO-Detection by indEXing (CODEX) Antibody Conjugation and Panel Creation

CODEX (now Phenocycler, Akoya Biosciences, UK) multiplexed imaging was executed according to the staining and imaging protocol previously described^23^. A panel of 56 antibodies were chosen to include targets that identify subtypes of the gastrointestinal epithelium, stromal cells, cells of the innate and adaptive immune system. Detailed panel information can be found in **Supplementary spreadsheet 2** and representative staining in **Figure S1**. Each antibody was conjugated to a unique oligonucleotide barcode, after which the tissues were stained with the antibody-oligonucleotide conjugates and validated so that staining patterns matched expected patterns already established for IHC within positive control tissues of the esophagus or tonsil. Similarly, Hematoxylin and Eosin morphology staining were used to confirm the location of marker staining. First, antibody-oligonucleotide conjugates were tested in low-plex fluorescence assays, and the signal-to-noise ratio was also evaluated at this step, then they were tested all together in a single CODEX multicycle.

### CODEX Multiplexed Imaging

The tissue arrays were then stained with the complete validated panel of CODEX antibodies and imaged^23^. Briefly, this entails cyclic stripping, annealing, and imaging of fluorescently labeled oligonucleotides complementary to the oligonucleotide on the conjugate. After validation of the antibody-oligonucleotide conjugate panel, a test CODEX multiplexed assay was run, during which the signal-to-noise ratio was again evaluated, and the optimal dilution, exposure time, and appropriate imaging cycle was evaluated for each conjugate **Supplementary data (Spreadsheet available upon peer review publication)**. Finally, each coverslip array underwent CODEX multiplexed imaging.

### CODEX Data Processing

Raw imaging data were then processed using the CODEX Uploader for image stitching, drift compensation, deconvolution, and cycle concatenation. Processed data were then segmented using CellSeg, a neural network R-CNN-based single-cell segmentation algorithm (https://github.com/michaellee1/CellSeg)^24^. After the upload, the images were again evaluated for specific signals: any markers that produced an untenable pattern or a low signal-to-noise ratio were excluded from the ensuing analysis. Uploaded images were visualized in Fiji/ImageJ (https://imagej.net/software/fiji/). For cell type subset classification, markers with a z-score above 1 were assigned as high expression, and those with a z-score below 1 were considered as low expression.

### Cell Type Analysis

Cell type identification was done following the methods developed previously^25^. Briefly, nucleated cells were selected by gating DRAQ5, Hoechst double-positive cells using CellEngine (https://cellengine.com/), followed by z-normalization of protein markers used for clustering (some phenotypic markers were not used in the unsupervised clustering). The data were over-clustered with X-shift^26^. Clusters were assigned a cell type based on average cluster protein expression and location within the image. Impure clusters were split or re-clustered following mapping back to the original fluorescent images.

To ensure consistent cell phenotyping across various disease states, cells from non-progressor, pre-progressor, and progressor datasets were processed as SingleCellExperiment^27^ objects. These datasets underwent batch correction using the Batchelor R package^28^ to achieve uniformity across samples. For dimension reduction and display, the uniform manifold approximation and projection (UMAP) analysis was performed utilizing the scater^29^ package, based on z-score normalized intensities. This method facilitated the evaluation of phenotypic consistency across the distinct experimental groups. A comprehensive analysis of the marker landscape within the phenotyped cell populations was conducted, elucidating the complex relationships between multiple markers. The proportion of each cell type and the mean signal intensity of individual markers were calculated and depicted using a dot plot (ggplot2^30^), offering an integrative perspective on marker expression across the different cell populations.

### Cell Type Quantification and Distribution

Cellular populations within each tissue core were quantified and normalized to the total cell count, encompassing both epithelial and stromal cells. The data are presented as the percentage of each cell type of the entire immune cell population. During spatial point pattern analysis, cells within the glandular space were isolated, with intra-epithelial lymphocytes identified as the immune component specific to this space. These lymphocytes were quantified and expressed as the number of immune cells per 100 glandular epithelial cells. Immune cells located outside of the Barrett’s glands were categorized as lamina propria immune cells. The distribution of these cell types across various stages of disease progression was illustrated using boxplots generated by the ggplot2^30^ package. To evaluate the independence of cell type distributions among non-progressor, pre-progressor, and progressor biopsy samples, a two-tailed Wilcoxon rank-sum test was conducted using the rstatix^31^ package, with statistical significance defined as p < 0.05.

### Immune Neighborhood Analysis

The immune neighborhood analysis was conducted following previously established protocols^32,33^. A systematic approach was employed, wherein a window comprising 10 cells was systematically chosen across the entire immune cell-type map of all cores. Each immune cell served as the central point (index cell) for its respective window during the analysis. Within each window, the number of each cell type among the nine nearest neighboring cells was calculated relative to the total number of cells within the window. The resulting data were organized in a matrix format, with each row representing an index cell and each column indicating the frequency of the corresponding cell type in the neighboring cells. To identify clusters of index cells with similar distributions of neighboring cell types, a mini-batch K-means clustering algorithm was employed. This method facilitated the grouping of index cells based on shared patterns of neighboring cell types, referred to as cellular neighborhoods (CNs). Following the clustering analysis, each index cell was assigned to the specific neighborhood to which it belonged.

### Spatial Point Patterns Analysis

The interactions between cell-cell and neighborhood-neighborhood were investigated using spatial point pattern analysis, which employed the x and y coordinates of various cell types or CNs as well as custom histological annotations. Ripley K cross function and cross-type Besag’s *L* function with an isotropic edge correction algorithm, provided by the spatstat^34^ package in R, were conducted on a per-TMA-core basis at various distances up to a maximum of 360 pixels (All). The analysis focused exclusively on immune cells located within the lamina propria. The cross-type function, which describes the relationship between two types of points, *i* and *j*, is defined as:

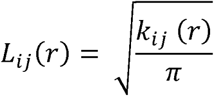

Where is the cross-type *K* function, which can be calculated as:

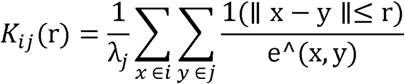

- r: The distance at which the function is evaluated.
- *L_ij_* (*r*) Cross-type Besag’s L function is a square-root transformation of the K function that linearizes the expected result under complete spatial randomness.
- *K_ij_* (*r*): The cross-type *K* function, which measures the expected number of type *j* points within a distance r of a randomly chosen type *i* point, normalized by the density of type *j* point.
- λ_j_: The density of type *j* points, defined as the number of points of type *j* per unit area.
- 1(||x−y||≤r): An indicator function that is equal to 1 if the distance |x−y| between points x (of type *i*) and y (of type *j*) is less than or equal to r, and 0 otherwise.
- e^ (x, y) The edge correction factor, which corrects for bias introduced by the edge of the observation window.

In a perfectly complete spatial randomness (CSR) point process, the *L* function should approximately equal the distance r, so the observed is plotted against the CSR as:

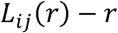

If the result for a given is greater than zero, it indicates that two-point types are more likely to be found in proximity, suggesting an attraction between them. Conversely, if the result is less than zero, it implies a tendency for these types to avoid each other, indicating repulsion at that scale. A result close to zero signifies a neutral relationship, with no significant attraction or repulsion observed.

### Interaction Score (IS) Calculation

The Diggle-Cressie-Loosmore-Ford (DCLF^35^) hypothesis test, as implemented in the spatstat^8^ R package, was utilized to determine whether the observed spatial pattern significantly deviated from CSR at the specified scale (distance) within eacy TMA core. This test was conducted at a 5% significance level, using the envelopes of L-functions derived from 39 simulated CSR realizations. Cores with P values ≤ 0.05 were considered significant, and the nature of interactions—whether attraction, repulsion, or random distribution was characterized within these significant cores.

Subsequently cell-to-cell and CN-to-CN interaction scores were then calculated by subtracting the number of repulsion cores from the number of attraction cores, then dividing by the total number of cores involved in cell-to-cell interaction or CN-to-CN interaction. This score was further corrected by multiplying it by the fraction of such present within the dataset. Comparisons among all cohort datasets were conducted and visualised by plotting matched interaction values for each cell-to-cell and CN-to-CN pair using ggplot2^12^. Statistical significance was determined using a Fisher’s exact test on the attraction, repulsion, and neutral confusion table between cohort datasets as implemented in the stats^17^ package. IS scores are detailed in supplemental spreadsheets available upon publication.

### Slide Preparation and Laser Capture Microdissection (LCM)

Serial sections, each 4μm thick, were prepared from CODEX Formalin-Fixed Paraffin-Embedded (FFPE) tissue samples. Carl Zeiss™ membrane slides (Carl Zeiss Microscopy GmbH, Germany, #415190-9042-000) were treated with ultraviolet (UV) light for 15 minutes to enhance adhesion before mounting the sections. To improve cell visualization, the slides were stained with a 0.5% methyl green and 0.5% Pyronin Y solution (Electron Microscopy Sciences, #26777-01) for 30 seconds to 1 minute.

For sequencing, three sections representing stromal regions and four sections representing epithelial regions were captured using P.A.L.M. laser capture microdissection (Zeiss) from a single core. These samples were collected in Adhesive Cap 500 opaque tubes (Carl Zeiss Microscopy GmbH, Germany, #415190-9201-000) and stored at −80°C until library construction. Cells from stromal regions underwent Smart-3SEQ^18^ RNA sequencing, while epithelial cells were analyzed by both Smart-3SEQ^18^ RNA sequencing and shallow-pass whole-genome sequencing.

### RNA Sequencing Library Construction

All RNAseq data will be made available upon publication. A 75 bp single-end RNA sequencing library was prepared following previously established protocols^36^ with slight modifications. In this study, the library construction process involved the utilization of 2uM of oligo dT and 50 μM of template switching oligo, which integrated 7-mer unique molecular identifiers (UMI). To initiate the process, the FFPE LCM lysis Mix was subjected to a 60°C incubation for 1 hour, followed by de-crosslinking at 70°C for 20 minutes. Reverse transcription was then performed under the following conditions: 42 °C for 30 minutes, 70 °C for 10 minutes, and a hold at 4 °C.

Subsequent PCR amplification was carried out using a G-storm (GS4) thermal cycler with the following program: an initial denaturation at 98 °C for 45 seconds, followed by 19 cycles of 98 °C for 15 seconds, 60 °C for 30 seconds, and 72 °C for 10 seconds, with a final extension at 72 °C for 60 seconds, and cooling step at 4 °C. The resulting library was purified with AMPure beads (Beckman Coulter, #A63880) at a 0.9:1 ratio of beads to sample. Library quantification was performed using the Agilent TapeStation (Agilent Technologies 4200) and D1000 ScreenTape (Agilent, #5067-5582).

The purified libraries were pooled and diluted to a final concentration of 4nM, and their size distribution was assessed within a range of 170–700 base pairs (bp). Single-end sequencing was performed using the Illumina NextSeq 2000 and NovaSeq platforms at the Genomics Facility of The Institute of Cancer Research, Sutton, UK in collaboration with Professor Trevor Graham.

### RNA Sequencing Analysis

Sequencing reads were assessed for quality using FastQC^37^ and trimmed for adapter sequences with the Trimmomatic^38^ package. UMI sequences were appended to read names, and adjacent GGG sequences were removed using an author-provided Python script. The processed reads were then aligned to the Genome Reference Consortium Human Build 38 with STAR^39^. PCR duplicates were removed and gene expression was quantified using UMI-tools^22^ and the Subread^40^ package.

To identify patterns associated with disease progression, genes with 5 or more UMI counts in at least 50% of the samples were selected and adjusted for sequencing platforms batch effects. The top 500 most variable genes were used for t-Distributed Stochastic Neighbor Embedding (Rtsne^41^) analysis. The x and y coordinates from the t-SNE reduction were clustered using k-means clustering (stat^42^) and visualized with a heatmap generated by ComplexHeatmap^25^.

Differential gene expression (DEG) analysis was performed with DESeq2^43^ to identify genes differentially expressed between clusters and disease progression statuses. DEGs with an average UMI count of at least 2 were depicted in volcano plots. Additionally, pathway analysis was conducted on the log2-transformed matrix from GSVA^44^ using parameters “kcdf = Gaussian” and “maxSize = 500”, utilizing hallmark gene sets from the Human Molecular Signatures Database^28^.

### Weighted Gene Co-Expression Network Analysis (WGCNA)

Weighted Gene Co-expression Network Analysis (WGCNA) was performed using the WGCNA^45^ R package. Initially, outlier samples were identified and excluded based on hierarchical clustering with Euclidean distance and average linkage. To ensure the robustness of the analysis, genes with fewer than 5 reads in at least 50% of the samples were removed. For network construction and module detection, the following parameters were used: a soft-thresholding power of 12, maxBlockSize = 5000, TOMType = “signed”, minModuleSize = 10, and mergeCutHeight = 0.25. These settings were optimized to enhance network connectivity and the sensitivity of module detection. The traits analysed are; RNAseq Kmeans clusters, non-progressors, pre-progressors and progressors, pathology (the pathology at time of progression) and time (time from sampling to time of progression or final endoscopy for non-progressors).

### Low-Pass Whole Genome Sequencing (LPWGS) Library Construction

Genomic DNA was extracted from LCM formalin-fixed paraffin-embedded tissue described above using the High Pure FFPET DNA Isolation Kit (Roche, #06650767001). The extracted DNA underwent repair with the NEBNext FFPE DNA Repair Mix (NEB, #M6330L), followed by bead-based purification and subsequent fragmentation at 37°C for 20 minutes. Dual indexing and the incorporation of P5 and P7 Illumina adapters were performed using NEBNext Multiplex Oligos (NEB, #E6609) in conjunction with the NEBNext Ultra II FS DNA Library Prep Kit (NEB, #E7805L) according to the manufacturer’s instructions. To ensure sufficient library fragment quantity for effective sequencing, each library was enriched through PCR amplification. The PCR cycling conditions included an initial denaturation at 98 °C for 30 seconds, followed by 14 cycles of 98 °C for 10 seconds, 65 °C for 75 seconds, and a final extension at 65 °C for 5 minutes, followed by a cooling step at 4 °C.

The resulting libraries were purified using AMPure beads (Beckman Coulter, #A63880) at a 0.9X bead-to-PCR reaction ratio and subsequently quantified with the Agilent TapeStation 4200 (Agilent Technologies) using D1000 ScreenTape (Agilent, #5067-5582). The purified libraries were then pooled and diluted to a final concentration of 4nM. Paired-end sequencing was conducted using the Illumina NextSeq platform at the Genomics Centre, Blizard Institute, Queen Mary University of London.

### LPWGS Analysis

DNA sequence reads were aligned to the human genome build 19 using the Burrows-Wheeler Alignment Tool (BWA^46^). Reads with a mapping quality score below 37 were excluded from subsequent analysis. PCR duplicates were identified, and base quality recalibration was carried out using the Genome Analysis Toolkit (GATK^47^). Chromosomal aberrations across the genome were quantified using the QDNAseq^48^ together with ACE^49^ R package with fixed-size bins of 100 kb.

### General Statistical Analysis and Visualization

All statistical analyses and graphical data visualizations were conducted using R (ver 4.1.2) and associated plugin software. Statistical analysis is described in each methodology.

**Resources:**

Anonymised META data file for preprint submission. All CODEX, RNAseq and sWGS data will be made publicly available upon peer-reviewed publication.

## Results

### The immune landscape of the NDBE mucosa changes and increases in diversity with progression to EAC

We assessed immune cell composition by performing Co-detection by indexing (CODEX) on pathologist-verified NDBE samples taken from non-progressor (never progressed, NP) (n=30 patients, 126 cores), prospective progressor (pre-progressor, PP) (n=5 patients, 50 cores) and progressor patients (already progressed, P) (n= 6 patients, 48 cores). A 56-plex antibody panel was used to profile as many cell types as possible (representative TMA CODEX staining **Figure S1**). Non-progressor NDBE samples were obtained from endoscopies from patients that did not subsequently develop EAC. Pre-progressor NDBE samples were obtained from endoscopies performed prior to the onset of dysplasia and progressor samples were taken from NDBE adjacent to adenocarcinoma. The workflow of our CODEX experiment is highlighted in **Figure 1A**.

**Figure 1.**
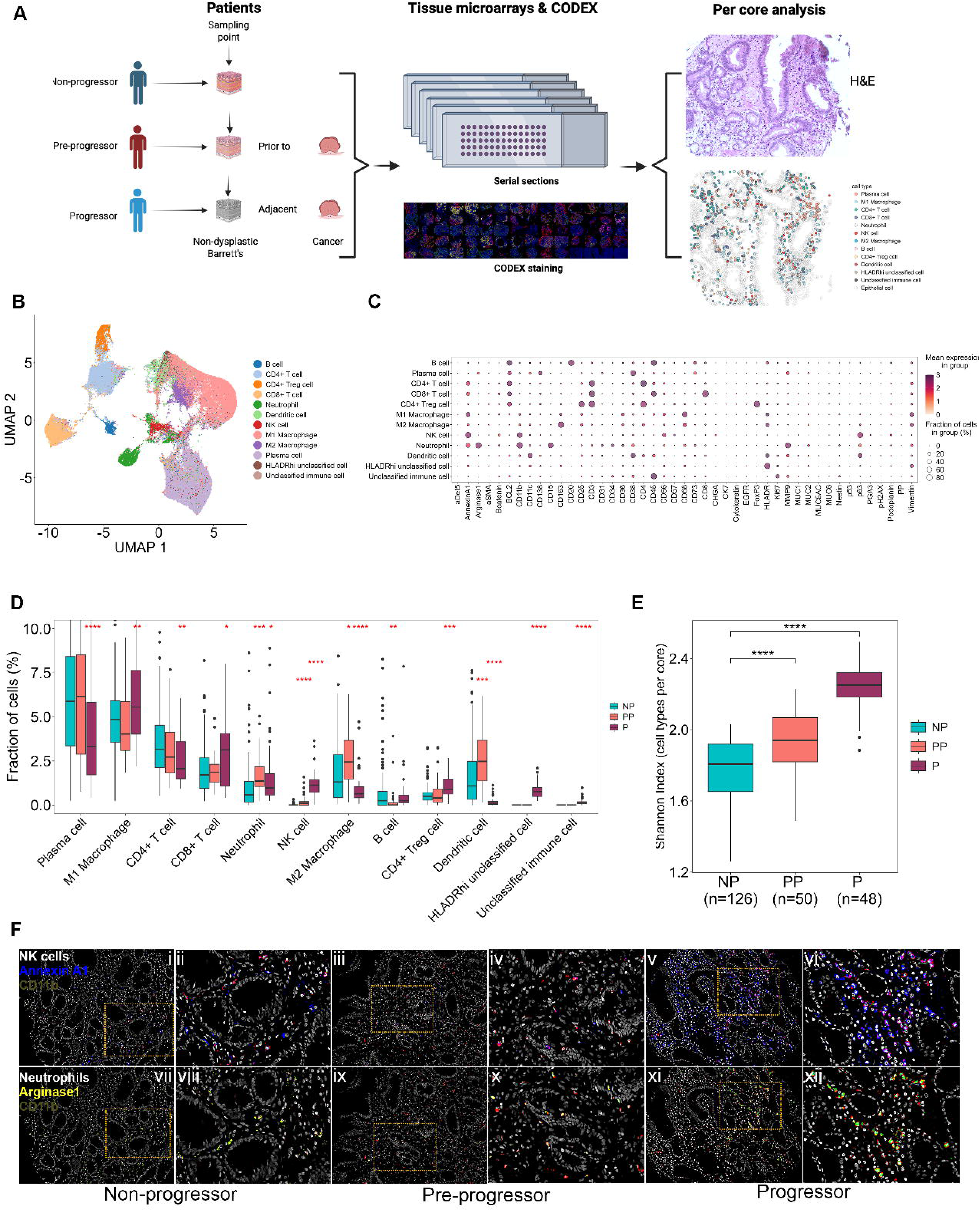
The metaplastic Barrett’s immune landscape before and during progression. (**A**)The workflow for CODEX analysis. Non-dysplastic Barrett’s esophagus (NDBE) samples are taken from *non-progressor (NP), pre-progressor (PP) and progressor (P)* patients and subjected to multiplex CODEX staining. Cells are mapped, quantified and their spatial relationship analysed on a per core basis. Created using Biorender. (**B**) UMAP dimensionality reduction illustrates all immune cell types present in all samples. (**C**) Dot plot of cell markers and their relative frequency and cell specificity from the CODEX panel. (**D**) Quantification for all identified immune cells as a percentage of all cells in NP, PP and P, respectively. (**E**) Shannon index of diversity of all immune cells in NP, PP and P core samples. (**F**) Representative CODEX images for NK cells (**i-iv**)(CD11b, red: AnnexinA1, blue. **i, iii, v** x100 magnification, **ii, iv, vi** x200 magnification. Green dotted box represents magnified area), neutrophils (**vii-xii**) (CD11b, red: Arginase1, green. **vii, ix, xi x100** magnification, **viii, x, xii** x200 magnification. Green dotted box represents magnified area) in non-progressors, pre-progressors and progressors respectively. (*P<0.05, **P<0.01, ***P<0.001, ****P<0.0001, Wilcoxon rank sum test).

We identified a total of 469,541 cells across all samples, and the diversity of immune cells is illustrated by UMAP dimensionality reduction analysis (**Figure 1B)**. Specifically, we identified 12 major immune cell types using 45 of the CODEX antibodies staining patterns (summarised in the progressor samples in **Figure 1C** and non-progressor and pre-progressor samples in **Figure S2, all cell maps for all cohorts are shown in Figures S3A-E**). Plasma cells and M1 macrophages were the predominant immune cell types observed in NDBE metaplasia across the three patient populations (**Figure 1D**). However, this comparative analysis showed that the dynamics of these immune cell types shifted in metaplasia as progression occurred. Thus, plasma cells (specifically CD138^+^ CD38^hi^ long-lived plasma cells), were reduced in progressor metaplasia compared with non-progressor metaplasia (**Figure S4A-D**). On the other hand, M1 macrophages increased and M2 macrophages decreased in progressors compared with non-progressors. Cytotoxic CD8^+^ T cells were also increased in progressor metaplasia. Notably, this was not reflected in pre-progressor samples that showed no significant differences in plasma cell nor in M1 macrophage populations (**Figure 1D**). Together, these data support the acquisition of a proinflammatory environment specific to progressor metaplasia.

Changes in the concentration of immunoregulatory cells were also identified. Immunoregulatory NK cells (CD3^-^, CD56^hi^, AnnexinA1^+^, CD11b^+^) and Arginase1^+^ neutrophils^50^ were increased in both pre-progressor and progressor metaplasia as illustrated by representative CODEX images of CD11b^+^ Annexin A1^+^ NK cells and Arginase1^+^ neutrophils across all NBDE categories (**Figure 1F)**. Additionally, the increase in neutrophils was associated alongside a switch in neutrophil MMP9^lo^ to MMP9^hi^ expression in progressors^51^, suggesting a conversion to a proinflammatory cell type upon progression (**Figure S5A-D**). Immunoregulatory CD4^+^ FOXP3^+^ Tregs were also increased in progressor metaplasia (**Figure 1D**). We also observed an increase in M2 macrophages and dendritic cells in pre-progressor metaplasia (**Figure 1D**). These data suggest the acquisition of an immunoregulatory response develops prior to a switch to a pro-inflammatory response in the metaplastic progression of BE.

We next probed the gene expression of the stromal microenvironment via laser capture microdissection (LCM) RNAseq. Investigation of immunoglobulin (Ig) expression led us to observe an increase in IgG4, IgM and IgA but not IgG1 antibody mRNA (**Figure S6A&B**) in both progressors and pre-progressors compared with non-progressors. Interestingly, gene expression of the classical complement component pathway C1A, C1B, C1C, C2 & C3 mRNA (**Figure S6C&D**) were also significantly increased in pre-progressor and progressor samples, a signature which is characteristic of macrophage interaction with Igs^15^. Overall, gene set enrichment analysis of LCM captured stromal samples revealed cell-specific gene expression associated with NK cells, neutrophils and dendritic cells (**Figure S6E-K**).

Based on these findings, we investigated the ecological diversity of immune cells within our data using the Shannon index (SI). All immune cell lineages were included in this analysis. We observed a progression-dependent increase in NDBE-associated immune cell diversity between all cohorts, illustrating an increase in immune cell complexity through metaplasia progression (**Figure 1E**). This is in concordance with our previous observation of NDBE gland diversity as a risk factor in progression of BE^52^. Overall, we observed that the immune microenvironment in NP to PP to P is specific to each stage and there appears a mixture of proinflammatory cells and immunoregulatory cells where both appear to increase sequentially with each cohort.

### Intra-epithelial immune cell composition is dysregulated in metaplastic progression

Intraepithelial immune (IEI) cells are a major immunosurveillance cell type in the gut. Most of these cells are CD8^+^ T cells but CD4^+^ T cells and sub-epithelial macrophages are also present^53^. To date, no study has examined the nature of IEI cells in BE, for which our muti-variable spatial data affords a unique opportunity. We thus quantified IEI cells by manually segmenting out the epithelial glands (**Figure 2A&B**). Magnified CODEX staining demonstrates that IEI CD8^+^ T cells and CD4^+^ T cells as well as CD68^+^ subepithelial macrophages can be detected (**Figure 2C**, magnified in **Figures 2D** respectively).

**Figure 2.**
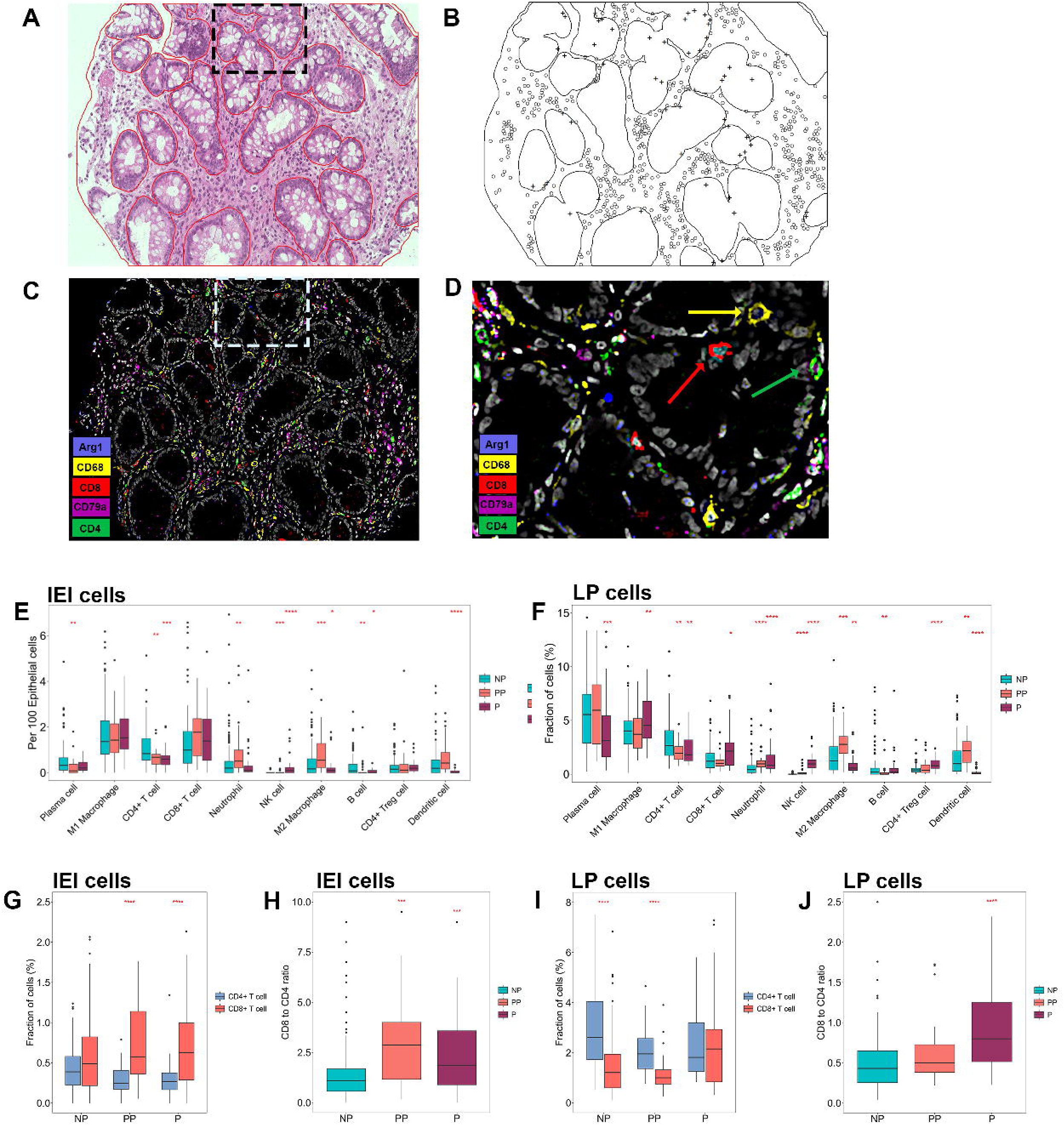
IEI cell and LP cell landscape in metaplastic progression. (**A**) H&E of a representative core (**B**) Map processed by Spatstat (cross=IEI cells, dot= LP cells) revealing the epithelial boundaries of (**A**). (**C**) 5-plex image of Arginase1, CD68, CD8, CD79a and CD4. (**D**) High power image of a high power glands revealing a CD8^+^ T IEI cell (*red, arrow*) and a sub-epithelial macrophage (*yellow, arrow*). (**E**) Quantification of IEI cells per 100 epithelial cells across NP, P and PP cores. (**F**) Quantification of lamina propria cells as a percentage of all segmented lamina propria cells. (**G**) Fraction of CD4^+^ and CD8^+^ T cells as percentage of IEI cells. (**H**) Ratio of CD8^+^:CD4^+^ T cell IEI cells. (**I**) Fraction of CD4^+^ and CD8^+^ LPLs as a percentage of all cells. (**J**) Ratio of CD8^+^:CD4^+^ T cell LP cells. ((*P<0.05, **P<0.01, ***P<0.001, ****P<0.0001, Wilcoxon rank sum test).

We observed a surprising variety of intraepithelial immune cells in all cores, the majority of these were CD8^+^ T cells and subepithelial M1 macrophages (**Figure 2E**). However, NK cells, plasma cells, dendritic cells, neutrophils and M2 macrophages were detectable in each cohort. Interestingly, we observed several differences in IEI cell concentrations in our three cohorts. There were significantly more neutrophils, M2 macrophages and dendritic cells and fewer CD4^+^ T cells in pre-progressor cores than in the non-progressors yet fewer M2 macrophages, dendritic cells and CD4^+^ T cells in the progressors (**Figures 2E**). Segmentation of IEI cells also permitted analysis of the lamina propria (LP) compartment immune cells, revealing a similar patterns of immune cell alterations (**Figure 2F**) compared to the bulk analysis (**Figure 1E**) indicating that IEI cells *per se* did not affect the overall analysis of bulk immune cells described earlier.

CD8^+^ T IEI cell concentrations were unaltered across all three cohorts; however, the ratio of CD8^+^:CD4^+^ T cells was significantly higher in pre-progressor and progressor NDBE cores compared with non-progressor NDBE cores (**Figure 2G&H**). We observed the opposite relationship when we compared the CD8^+^:CD4^+^ ratio among the LP cells where the ratio of CD8^+^ T cells to CD4^+^ T cells was lower in non-progressor and pre-progressors compared with progressor NDBE (**Figure 2I&J**). There is a clear dichotomous relationship between the numbers of CD8^+^ T cells and CD4^+^ T cells in the metaplastic epithelium and this appears to have important implications for progression to cancer in BE. Overall, IEI cells are a rich population of BE immune cells with distinct differences between non-progressors, pre-progressor and progressor metaplastic samples suggesting dysregulation of immune surveillance within the BE epithelium.

### Disordered immune cell interactions in pre-progressor and progressor NDBE samples

In view of the altered frequency of immune cells with progression, we next examined pairwise spatial relationships among immune cell types as indicator of functional interaction with each other, either positively or negatively. We used Ripley K cross type function permutations to determine pairwise cell interactions (illustrated in **Figure 3A-C** and described in the Methods). To determine which immune cell pairs that showed attraction to or repulsion from each other, we calculated Ripley L values (transformed from K values) over all distance from each index cell to their comparator cells and plotted these against a complete spatial randomness null-model (CSR, simulated 39 times). An L value greater than CSR indicates pairwise attraction (e.g., CD4^+^ T cells and plasma cells (**Figure 3D**)), while a value below indicates repulsion (e.g., CD4^+^ Tregs and dendritic cells (DCs, **Figure 3E**)) and any value that lies within the minimum and maximum CSR values represented a random association. An example of Ripley L value plots across each condition comparing plasma cells and M1 macrophages is shown in **Figures 3F-I**. Additional examples are shown in **Figure S7** Interaction scores were then compared in a pairwise fashion between non- and pre-progressors (**Figure 3J**, M1 macrophage interactions highlighted in green, NK cell interactions highlighted in blue), non- and progressors (**Figure 3K**) and pre- and progressors (**Figure 3L**) at 100 arbitrary distance units. A summary dot plot of all cell-pair interactions is shown in **Figure 3M**. Similar data was obtained from other distances and are shown in **Figure S8**.

**Figure 3.**
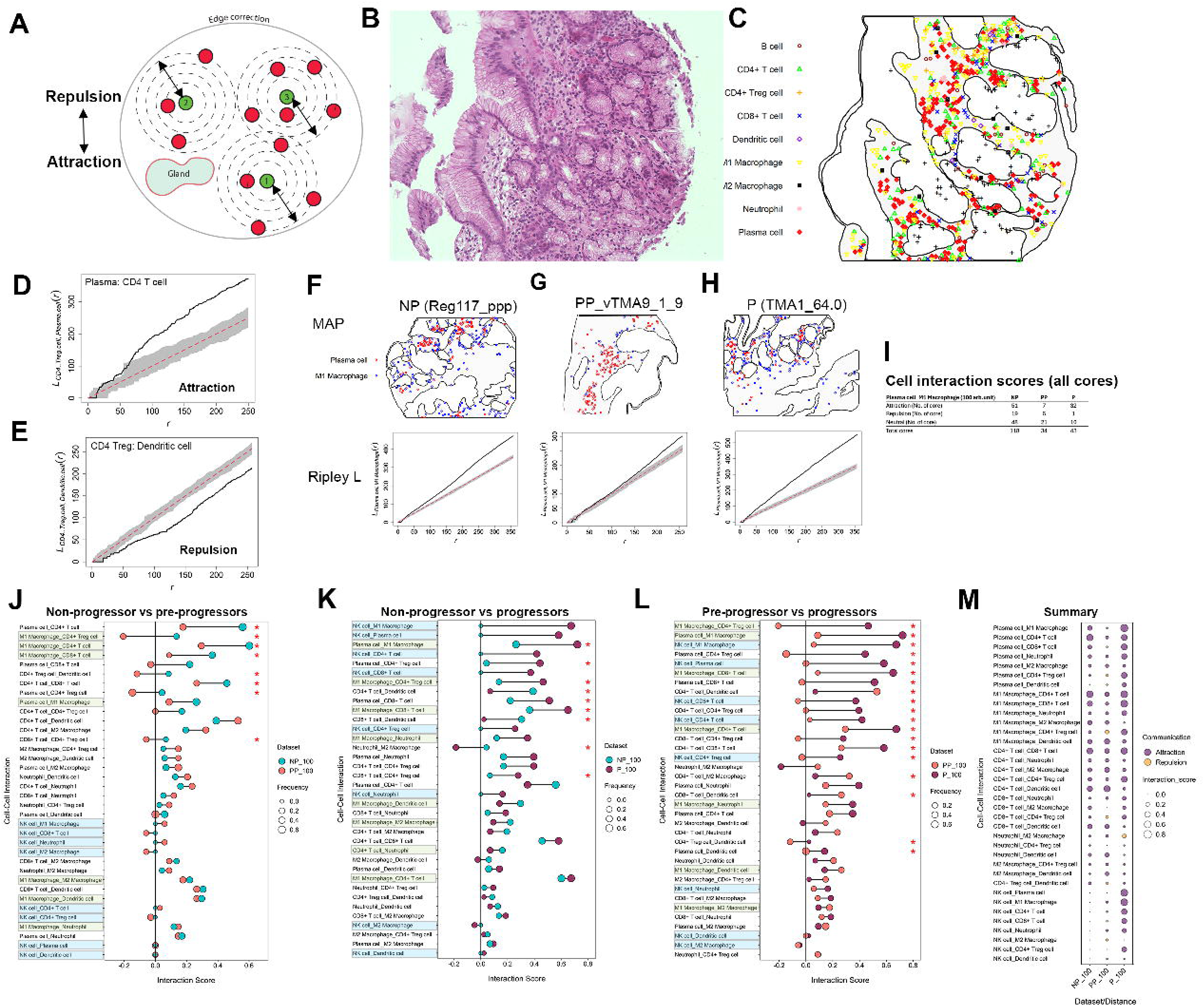
Cell to cell interactions in metaplastic progression. (**A**) Illustration of the Ripley K function analysis of attraction and repulsion (arrows represent distance) on pairwise cell comparisons (Green and red hypothetical cells). Concentrations of red cells are quantified at increasing radii from each green cell. (**B**) Representative core H&E. **(C**) Map highlighting center pixel of all immune cells within the representative core, epithelial cells are excluded. (**D**) An example of cell-to-cell attraction (Plasma cell: CD4 T cell). (**E**) An example of cell-to-cell repulsion CD4 Treg: Dendritic cell. (**F**) TOP: map of plasma cell and M1 macrophages in a non-progressor core. BOTTOM: with Ripley L values plotted against complete spatial randomness (CSR) showing a strong attraction. (**G**) In a pre-progressor showing weak distant attraction. (**H**) In a progressor showing strong attraction. (**I**) The number of cores exhibiting attraction, repulsion or neutral (random) distribution in each condition. (**J**) Illustration of differential mean interaction scores between non- and pre-progressors. The Size of each circle represents the frequency of cell pairs deviating from CSR. Highlighted are M1 macrophage (green) and NK cell (blue) interaction score comparisons. (**K**) Interaction scores for non- and progressor samples.. (**L**) Interaction scores for pre-to progressor samples. (**M**) Summary dot plot of interactions scores at 100 distance units. Size of circles represent mean interaction scores (*P<0.05, Fisher’s exact test).

It appeared from the observed interaction score values that in non-progressor patients there was a strong spatial attraction between M1 macrophages and plasma cells, T cells and DCs (**Figure 3J)**. However, this spatial cellular organization was lost in pre-progressor samples where most immune cell attractions drifted to a random distribution (**Figure 3J**). This is in stark contrast with non-vs progressor scores (**Figure 3K**) that show a significantly stronger attraction between M1 macrophages and plasma cells despite progressors exhibiting a decrease of plasma cells.

NK cells were rare in non-progressor patients but increased significantly in pre-progressor patients (**Figure 1D**. However, most NK cell interactions with other immune cell types in these pre-progressor patients appear randomly distributed (**Figure 3J**). This dramatically changes in progressor NDBE where a strong attraction between NK cells and plasma cells, M1 macrophages, CD4^+^ T cells and CD8^+^ T cells was observed (**Figure 3L**). Conversely, NK cells did not interact with many of the immunoregulatory cells (FoxP3^+^ Tregs, M2 macrophages) and this was unchanged from pre-progressors to progressor samples. With regards to neutrophils, despite their increase in cell number in progression, we did not observe any significant interactions with any other immune cell type, suggesting that their function was either non-specific (MMP9^lo^) or promoted angiogenesis (MMP9^hi^)^51^. Overall, the emergence of multiple cell interactions within the disorganized state of pre-progressor as lesions progressed indicate the emergence of a complex coordinated pro-tumorigenic the immune response. **Figure 3M** shows a summary dot plot of all attraction and repulsions between each cell pair and the NDBE progression status.

### Cellular organizations are altered before and after metaplastic progression

To examine higher-order spatial relationships given the multiple pairwise immune cell interactions and to expose potential multi-cellular functional units in the tissue, we investigated whether specific cell neighborhoods^32^ (CNs) existed within NDBE and how these were altered between pre-progressor and progressor metaplasia. Unlike prior work on CN, here we excluded all non-immune cells^21^. This enabled us to investigate immune cell neighborhood architectures irrespective of other cell type enrichments. CNs were also identified through combining all three patient data sets then performing clustering using a mini-batch K-means clustering algorithm (see methods) to enable comparison of common CNs.

A total of eight CNs were identified (**Figure 4A**) that represented CNs with either a high density of single cell types or those that were more mixed. We observed regional clusters of CNs from the same neighborhood as we would expect based on local microenvironments (**Figure 4B-G**). **Figure 4H** illustrates the cellular component characteristics of all identified CNs all three NDBE progression cohorts. Two major plasma cell CNs (CN3 & CN7) were identified with the first (CN7) almost exclusively composed of plasma cells. Plasma 2 CN (CN3) was also composed primarily of plasma cells but also contained CD4^+^ T cells and M1 macrophages. One CN predominantly consisted of M1 macrophages (CN4) while another macrophage-enriched CN (CN0) consisted of an equivalent number of M1 macrophages and plasma cells. Three complex CNs were identified that either contained mainly dendritic cells (CN1), CD4^+^ T cells (CN2) or M2 macrophages (CN6), respectively. A CN that was almost entirely composed of B cells (CN5) was also detected but only a few cores presented with this CN. While the cell composition of CNs was consistent across NDBE progression, NK cells and neutrophils appeared to be present in nearly all pre- and progressor CNs with their frequency increasing in a linear manner across progression. This suggested that NK cell and neutrophil presence was independent of CNs. The cellular complexity of CNs also appears altered with progression (**Figure S9**). We observed an increased Shannon diversity in CN1 and CN6 in pre-progressor samples, and an increase in CN0, CN2, CN4 and CN6 in progressor samples suggesting that the changes in cellular diversity observed in **Figure 1E** are tied to specific neighborhoods. **Figures S10A-E** show the individual core neighborhood maps for all samples.

**Figure 4.**
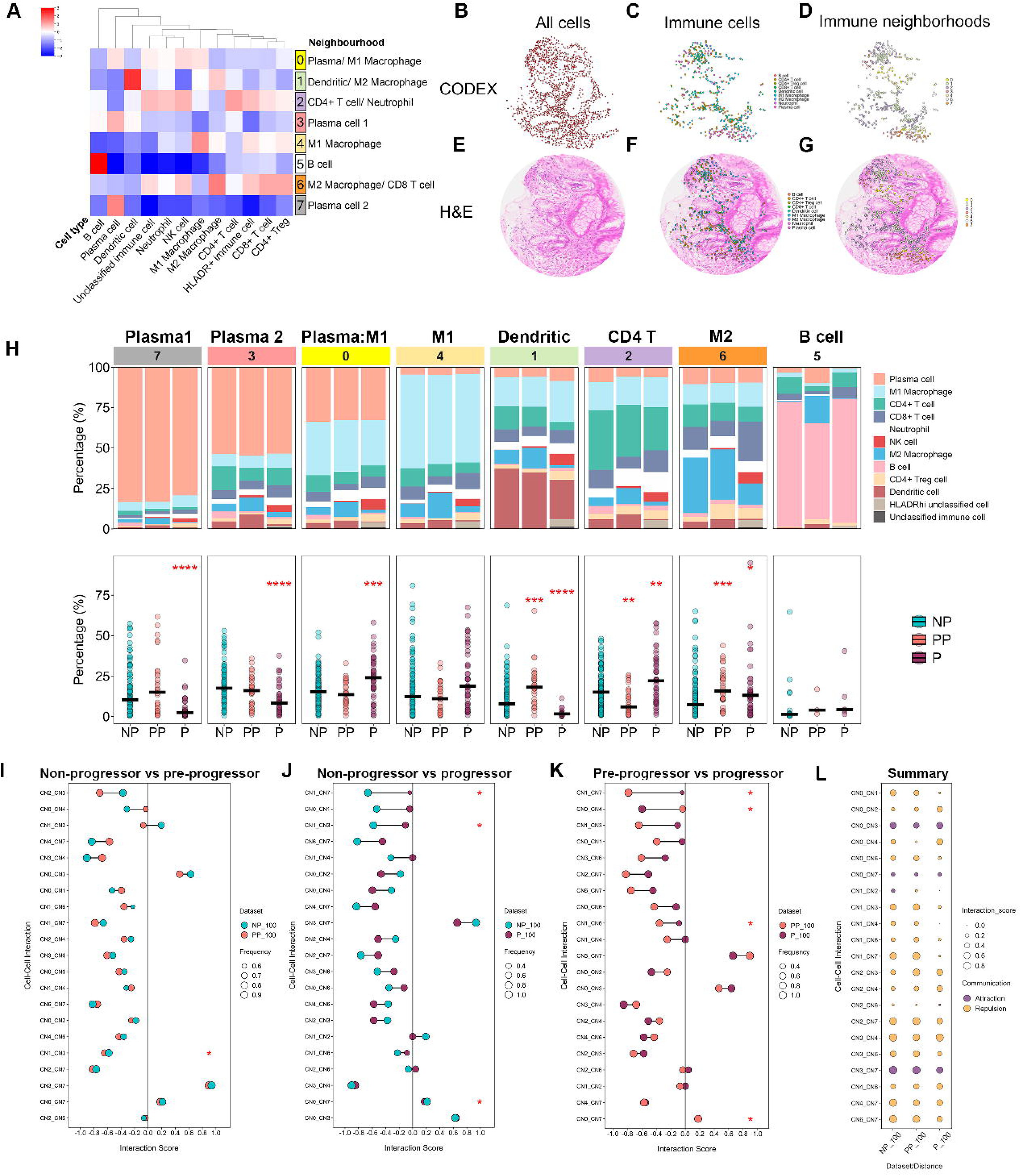
Cell neighborhoods in metaplastic progression. (**A**) Heatmap of nearest neighbor clustering across all immune cells and all samples. Neighborhoods named after dominant cell type(s). (**B-D**) All cells, neighborhoods and individual immune cell types as per x, y coordinates. (**E-G**) Mapped onto the respective H&E image taken after CODEX completion. (**H**) TOP: Cell composition of CNs (% of total immune cells) in order of non-progressor (NP), pre-progressor (PP) and progressor (P) samples. BOTTOM: Percentage of cells representing each neighborhood in NP, PP and P samples. Pairwise neighborhood-to-neighborhood in (**I**) Non-to preprogressors, (**J**) Non-to progressor and (**K**) Pre-to progressor samples. (**M**) Summary of overall neighborhood interaction scores across all samples. (*P<0.05, **P<0.01, ***P<0.001, ****P<0.0001, Fisher’s exact test).

We observed significant alterations in CN frequency and composition as metaplasia progressed (**Figure 4I**). Pre-progressors exhibited marked reduction in CD4^+^ T cell CN (CN2) and an increase in the number of cores containing M2 macrophages/CD8^+^ T cells (CN6) and dendritic cell CNs (CN1) compared with non-progressor CNs respectively. Progressor CNs appeared most altered compared with non-progressor CNs. A decrease of plasma 1 (CN3) and plasma 2 (CN7) CN correlated with an overall decrease of plasma cells in progressor samples described above suggesting that plasma cells, in spatial isolation from other cells, were lost. An increase in the proportion of CN0 plasma cell/ M1 macrophage neighborhood was observed in progressors compared to non-progressors. Plasm cells contained within CN0 were predominantly CD38^hi^ CD138^+^ (long lived) (**Figures S11A & B**). This suggests that while overall numbers of plasma cells decreased as the lesion progressed, those that remained were highly active (likely producing Ig mRNA, **Figure S6**).

Interestingly, despite an overall reduction in CD4^+^ T cells among progressor patients, the CD4^+^ T cell-enriched CN (CN2) was found to be increased (**Figure 4H**). This CN included a wide range of cell types, suggesting a more heterogeneous immune microenvironment in progressors. In contrast, the CD8^+^ T cell-enriched CN (CN6) appeared to mirror the elevated presence of CD8+ T cells in the stromal regions of progressor samples. Notably, the ratio of CD8^+^ to CD4^+^ T cell-enriched CNs (CN2 and CN6, **Figure 4H**) corresponded closely to the relative abundance of these cells in the epithelium (**Figure 2G&H**). Together, these observations suggest that the cellular composition of CNs—especially those involving plasma cells and M1 macrophages—undergoes marked remodelling in NDBE as patients progress toward EAC.

Previous studies have shown a spatial relationship between distinct CNs in EAC epithelium ^21,25^. Thus, we developed multiple analytical methods to examine these relationships, such as permutation testing and neighborhoods of neighborhoods^21,33^. In a similar fashion, to explore whether CN-CN relationships are affected by NDBE progression, we applied Ripley’s K function analysis to assess the spatial pairwise association of CNs in all samples (**Figure 4I-K**). This enables investigation of CNs that may share borders.

While most CNs exist in isolation from each other, which is expected because they are likely self-contained neighborhoods, plasma cell CNs (CN0, 3 & 7) exhibited significant spatial clustering together across all 3 cohorts. Interestingly, we observed no significant alterations in interactions between CNs in non-progressor and pre-progressor samples (**Figure 4I**). However, in progressor samples we observed a significant shift from isolated CNs towards a more random distribution (**Figures 4J&K**). A loss of organized CNs in progressor samples and the changes in immune cell distributions, coupled with an increase in cellular diversity suggests that the pro-oncogenic immune environment becomes more disorganized, with many cell types interacting with each other at increased levels in pre- and progressors compared to non-progressors. The cell neighborhood interactions are summarized in **Figure 4L**.

### Gene expression analysis reveals pathways associated with progression in NDBE

To investigate the molecular underpinning of the stroma in NDBE from all cohorts, we performed LCM microdissection RNAseq on multiple stromal areas identified in each core (illustrated in **Figure 5A-D**). Dimensionality reduction tSNE maps did not reveal robust clusters associated with progression status (**Figure 5E**) but revealed three distinct clusters of gene expression in k-means clustering (**Figure 5F**). Frequency analysis of each individual library showed that there was a significant enrichment for pre-progressor and progressor patients in cluster Kmeans1(K1) compared with clusters K2 and K3 (**Figure 5G**). Each cluster is illustrated as a supervised heatmap of the 500 most variable genes (**Figure 5H**). GSVA analysis (**Figure 5I**) revealed that K1 showed highly enriched gene expression in multiple pathways, including inflammatory (IFNγ response, TNFα signaling, inflammatory response and IL-2/Stat5 signaling Hallmark gene set pathways) and cancer-related (EMT, KRAS signaling, P53 pathway Hallmark gene set pathways). By contrast, cluster K2 exhibited intermediate levels of gene expression, and cluster K3 showed minimal or ‘cold’ expression for these pathways.

**Figure 5.**
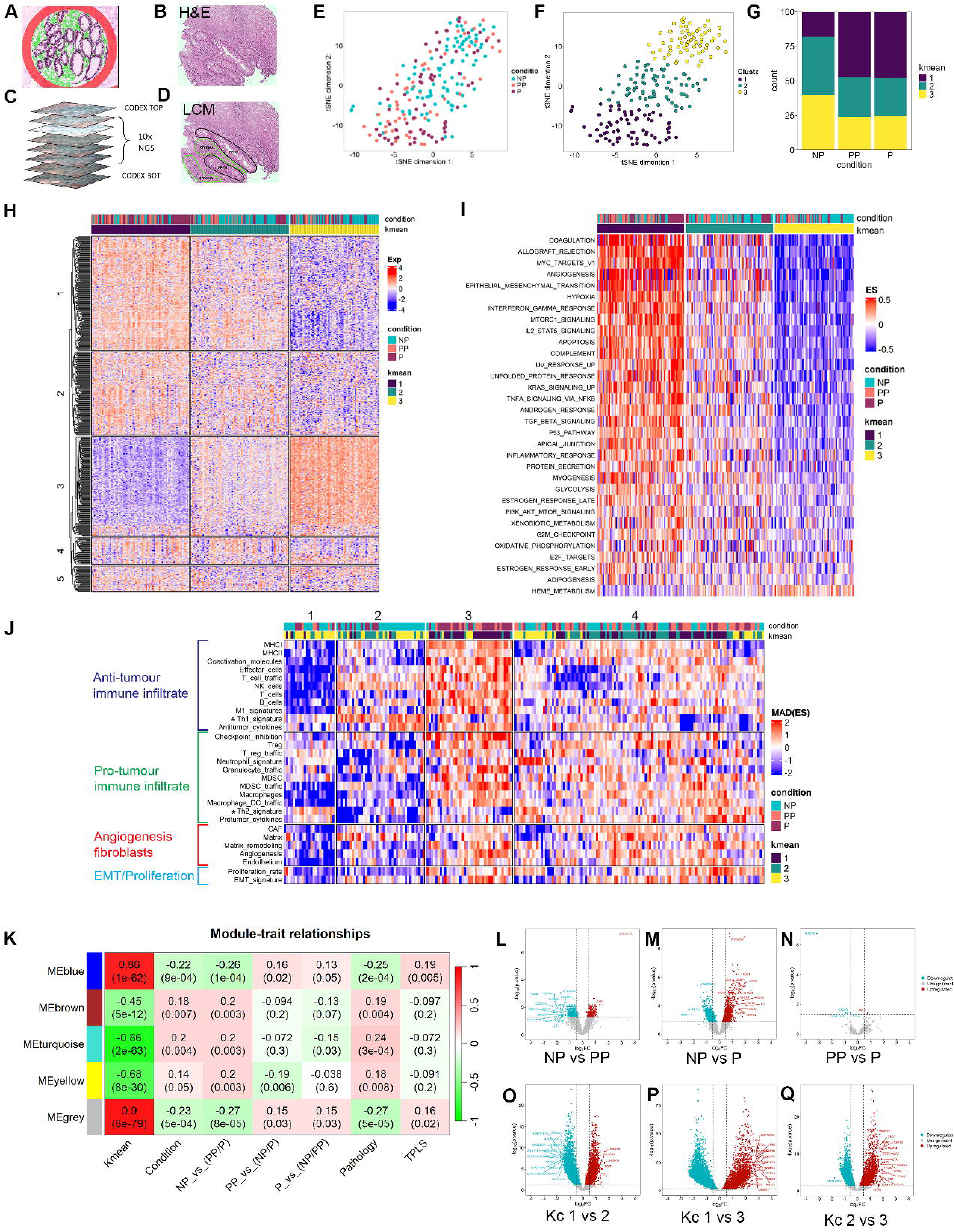
Gene expression analysis of metaplastic progression. (**A**) RNAseq protocol showing stromal (green) and epithelial (purple) regions of interest. NGS LCM slides are cut between two CODEX slides (top and bottom). (**B**) H&E image of the studied core. (**C**) NGS LCM slides are cut between two CODEX slides (top and bottom). (**D**) Area of interest for laser capture microdissection: glandular epithelium (black), stroma (green). (**E&F**) t-distributed Stochastic Neighbor Embedding (t-SNE) demonstrating K means clusters of gene expression. (**G**) Percentage of non-progressor (NP), pre-progressor (PP) and progressor (P) samples associated with each K means cluster. (**E**) Heatmap of the 500 most differentially expressed genes supervised for K means cluster. Condition = NP, PP and P. **(I**) GSVA analysis of differentially expressed pathways supervised for K means. (**J**) Bagaev et al.^54^, devised a pan-cancer microenvironment gene expression signature that was successful in predicting immunotherapy outcome based on 4 expression signatures of anti-tumor, pro-tumor, angiogenesis/fibroblasts and EMT/proliferation. Based on this report, we subjected our LMC RNAseq dataset to approach an unsupervised analysis in NP, PP and P and K means clusters. (**H**) WCGNA analysis for Kmeans, NP, PP, P, pathology histology at progression and time (TPLS, time to progression or time to last known surveillance). (L-Q) Volcano plots demonstrating >log2 fold expression and adj P value<0.05 for (**L**) NP vs PP.), (**M**) NP vs P, (**N**) P vs P. Comparison of Kmeans clusters (**O**) 1 vs 2, (**P**) 1 vs 3 and (**Q**) 2 vs 3.

Applying the approach described in Bagaev et al.^54^(**Figure 5J**), we observed an immune cold cluster (cluster 1) enriched for non-progressors and K3 samples that expressed a Th2 cytokine signature. An additional cluster (cluster 2) also enriched for non-progressors/K3 samples showed an immune cold environment but also a high Th1 signature. Pre- and progressor/K1 samples were enriched in an immune hot cluster that lost Th2 cytokines but expressed Th1 cytokines (cluster 3). A fourth cluster (4) was evenly distributed across all 3 patient cohorts. Interestingly, K1 was associated with both an anti- and pro-immune signatures, providing further evidence for a heterogenous immune response with components that appear to counterbalance each other.

To better understand the relationship between gene expression changes and progression, we performed weighted correlation network analysis (WCGNA), linking gene expression patterns with specific traits such as K means, progression status, future pathology (pathology at final known timepoint) and time to last known outcome (e.g., at last endoscopy). **Figure 5K** confirmed the strong correlation with K means clustering (illustrated as MeBlue and MeGrey); while a weaker, yet significant correlation with progression was observed (illustrated as MeBrown, MeTurqoise and MeYellow). Future pathology (that observed at the time point of progression or last endoscopy) also showed a weak, but significant association with endpoint histology similar to progression status, but no clear differences between endpoint pathology could be discerned. Biopsy time (TPLS) before endpoint did not show any relationship.

Individual gene expression changes across the transcriptome were visualized by volcano plots for both progression status and the K means clusters. Interestingly, we observed a large group of differentially expressed genes between non-progressors and pre-progressors (**Figure 5L**) and, also between non-progressors and progressors (**Figure 5M**) that concords with the pathways described in **Figures 5I&J**. Unexpectedly, comparing pre-progressors and progressors yielded almost no significantly differentially expressed genes (**Figure 5N**). Volcano plots confirmed the significant gene expression changes between the K means clustered (**Figures 5O-Q**). In summary, the stromal gene expression profiles of pre-and progressor samples were very similar, in that the stroma of pre-progressors already reflected a progressor state, an important observation noted in our previous publication^21^.

### Potential role for other stromal cells in regulating the BE microenvironment

Analysis of stromal signatures using WCGNA (**Figure 5K**) revealed that one module (MeYellow) was particularly enriched for extracellular matrix genes in pre-progressor samples. These were confirmed on an individual gene basis (**Figure 6A**), showing that collagen genes, MMP2, LUM and SPON2 levels were significantly increased in pre-progressor compared with non-progressor samples. Interestingly, only some of these (MMP2, LUM, COL6A3 & COL5A2) were also increased in progressor metaplasia. ‘Enrichr’ analysis confirmed that these genes were likely expressed by fibroblasts (**Figure 6B**).

**Figure 6.**
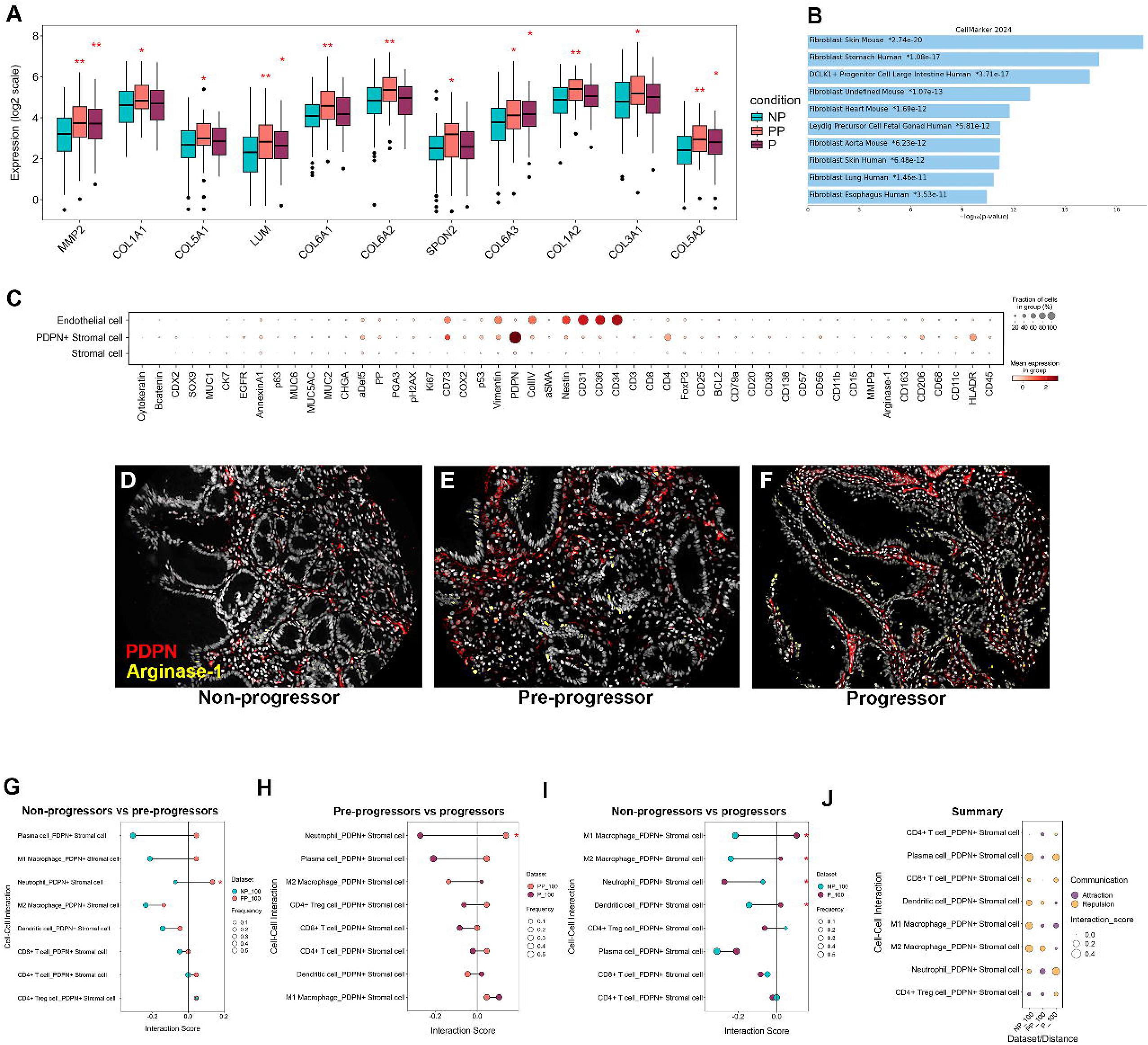
Fibroblast interactions and gene expression alters through progression. (**A**) WCGNA yellow module, altered in pre-progressors is enriched for extracellular matrix genes. (**B**) Enrichr analysis of cell lineage from yellow module. (**C**) Dot plot of cell specific CODEX panel markers for non-immune cells. CODEX staining images for PDPN and Arginase1 for (**D**) non-progressor, (**E**) pre-progressor and (**F**) progressor metaplasia. Pairwise, cell interaction scores at 100 arbitrary distance units for (**G**) non-progressor: pre-progressor, (**H**) pre-progressor: progressor and (**I**) non-progressor: progressor samples. (**J**) A summary dot-plot for all conditions showing the strength of attraction and repulsion interactions. (*P<0.05, Fischer’s exact test).

Because we saw an upregulation of genes from other stromal cells besides immune cells, we investigated other stromal cell types in our CODEX dataset. Dot plot for pre-progressor samples identified highly expressed non-immune stromal cell types including PDPN^+^ stromal fibroblasts, SMA^+^ smooth muscle cells and CD31^+^ CD34^+^ CD36^+^ endothelial cells (**Figure 6C**, representative CODEX images for PDPN^+^_Arginase1^+^ cells are shown in **Figure 6D-F**). While data presented earlier (**Figure 3D**) showed only random spatial distributions across immune cells in pre-progressors, the PDPN+ cells exhibited instead attraction towards neutrophils only in pre-progressor NDBE (**Figure 6G-J**). These data provided a potential explanation for how increase in MMP9 occurred in neutrophils, although we cannot ascertain that PDPN+ cells are exclusively fibroblasts from our dataset.

### Copy number variations (CNVs) of metaplastic epithelial cells correlate with specific immune cell distributions

Our analysis suggests the important role of immune cell composition and organization with progression. Therefore, we hypothesized that we should also identify relationships between immune cell composition and epithelial copy number variations which have been reported to exhibit a strong association with progression to dysplasia^21^. Indeed, progression to dysplasia or cancer can be affected by mutations or copy numbers affecting *TP53*^12,13,55^ and/or a loss of *CDKN2A* (p16)^56^ ^57^but the relationship between immune cells with epithelial genomic alterations in progressive BE remains poorly understood. We therefore performed shallow pass whole genome sequencing (sWGS) on LCM-isolated epithelium from serial sections taken from the cores subjected to CODEX (**Figure S12A-H** for representative sWGS plots) and then correlated copy number changes with immune cell frequency in an unsupervised fashion. We did not expect to encompass the full spectrum of CNVs previously reported^7^ but rather to credential the association of pro-tumorigenic immune cell profiles in stroma adjacent to epithelial lesions bearing such CNVs in our cohort(s). Two major clusters were identified that strongly separated progressor and non-progressor NDBE (**Figure 7A**). Samples displaying diploid genomes were most common. However, chr18p amplicons, chr9 deletions and p53 loss (determined by sWGS and CODEX respectively) were observed. Chr18 amplicons were detected exclusively in progressor epithelium and chr9 deletions were almost always detected in non-progressor epithelium (**Figure 7A**). p53 expression loss as measured by CODEX, strongly associated with progressor samples; conversely, one non-progressor showed high expression of p53 (**Figure 7A**). Additionally, one progressor case had both a chr18 amplicon and a chr9 deletion. Other cases contained CNVs that were restricted to a single sample, such cases being grouped together as ‘CNVs’.

**Figure 7.**
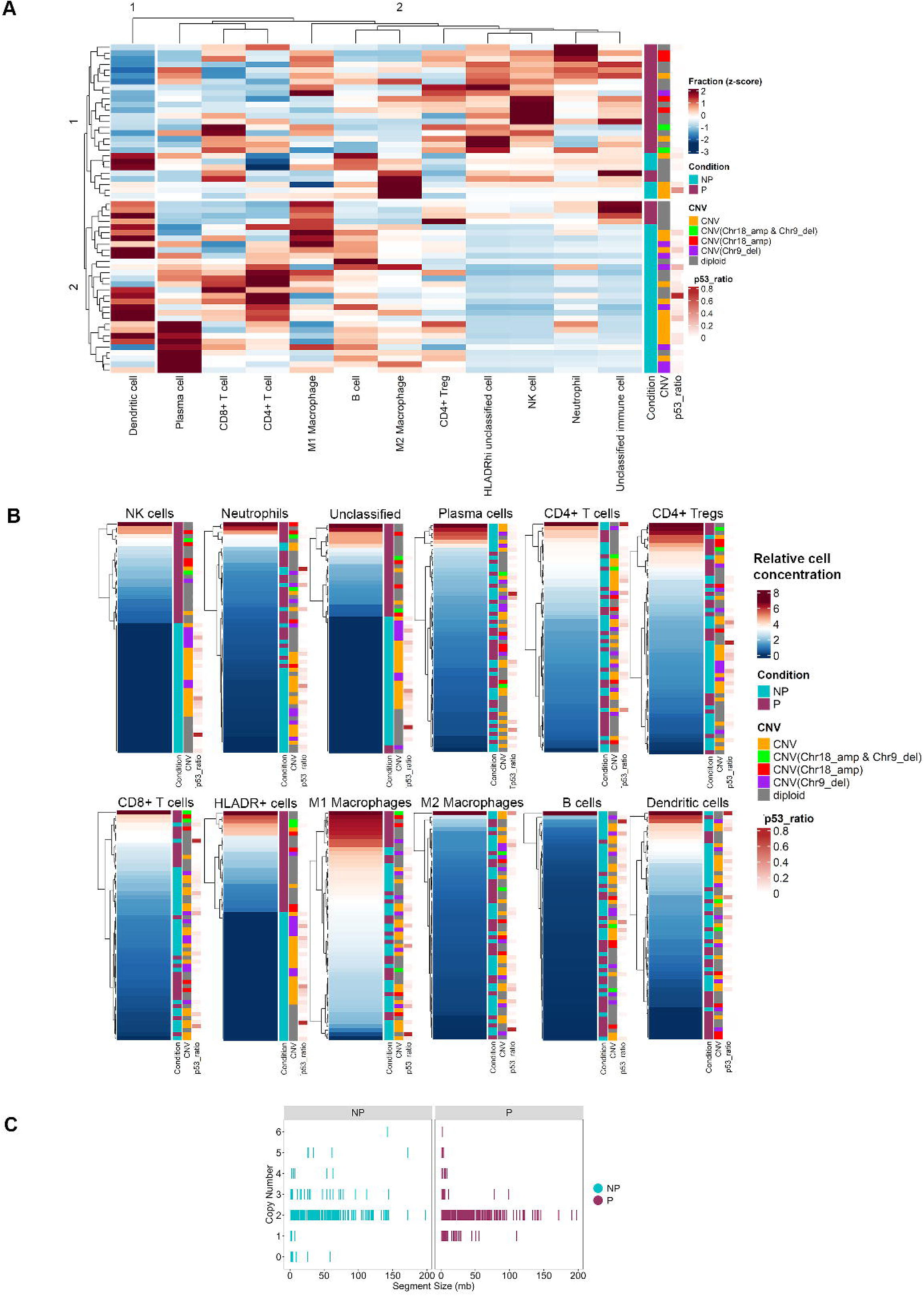
Immune cell correlations with copy number variations. (**A**) Unsupervised heat map of immune cell expression correlation with non-progressors or progressors (NP or P) with copy number changes (Orange= all CNVs except in chromosome 18 and 9, Green=Chr18 amplification and Chr9 loss, Red= Chr18 amplification only, Purple= Chr9 deletion only, Grey= diploid and Gradient= P53 expression level). (**B**) Correlation with NDBE progression status and CNVs supervised with concentration of each individual immune cell type. (**C**) Frequency of segment sizes of identified CNVs within non-progressor(NP) and progressor (P) NDBE.

Unsupervised clustering of cellular expression with CNVs showed a strong correlation between chr18 amplifications and enrichment in NK cells and neutrophils, whereas such correlations were not observed when contrasting CNV-bearing epithelium with M1 macrophage, plasma cell, CD4^+^ or CD8^+^ T cell profiles (**Figure 7B**). Moreover, there was a correlation between NK and neutrophil cell frequency and other CNVs but this association was not seen for other immune cells except for rare, uncharacterized HLADR^+^ or CD45^+^ lin-cells that we could not further characterize from our CODEX panel. This does not discount the possibility that correlation of CNVs and immune cell changes arise independently of each other. **Figure 7C** shows the frequency of all CNVs and their respective genomic segment sizes showing no apparent difference. From these data, we infer that NK cells and neutrophils are strongly associated with chr18 amplifications and M1 and plasma cells are likely to be independent of chr18 amplification because they are frequent in the non-progressor NBDE. These data suggest that there is a strong relationship between specific immune cells and the generation of specific epithelial CNVs in progressive BE.

## Discussion

This study highlights significant changes in the spatial organization and immune cell composition of the premalignant metaplastic stroma and how this integrates with gene expression prior and during carcinogenesis. Particularly, we investigated how the immune landscape in Barrett’s esophagus metaplastic lesions develops during this process.

Gaining a detailed understanding of the alterations of cellular interactions and organization across BE will eventually improve our ability to predict which BE patients are at higher risk of progressing to EAC. To begin to understand this process defining the microenvironment that associates with the progressor NDBE is critical^5^. To date, studies on pre-progressor NDBE have focused on the genomic landscape which has revealed altered genotypes in patients, years before progression^7,8,10,22^. Here, we are the first study to investigate how the immune response changes in NDBE as the risk of cancer progression increases.

One of our most surprising findings is that there is an equilibrium between pro-inflammatory and counter immunoregulatory programs in the context of NDBE, which is altered in the evolution of non-progressor to pre-progressor to progressor states. The non-progressor immune landscape is rich in M2 immunomodulatory macrophages, plasma cells and dendritic cells that appear to show a strong reciprocal attraction to one another and maintain specific plasma cell-enriched neighborhoods. The pre-progressor stroma is characterized by an increased number of immunoregulatory cells such as M2 macrophages and immunoregulatory NK cells, which contrasts with an increase in neutrophils. We observed a loss of immune organization in pre-progressor cell interactions with a significant drift to random spatial distribution between M1 macrophages and most other immune cells except for dendritic cells which maintain interactions with M1 macrophages and CD4+ T cells. These changes appear in the setting of increased collagen and MMP2 gene expression. The progressive NDBE displays a further significant increase in regulatory NK cells, countered by a concomitant increase in pro-inflammatory M1 macrophages and neutrophils. Interestingly, there is a switch to an immune hot environment where M1 macrophages strongly interact with most other immune cells. This is counteracted by regulatory NK cells, despite being a minority population in progressive NDBE, as these NK cells are strongly attracted to most other immune cells with the exception of neutrophils. The diversity of the immune repertoire in BE may reflect the complex interplay between pro-inflammatory signaling driven initially by acidic reflux and subsequent anti-inflammatory responses by the BE epithelium and stroma. Cellular diversity is a known cancer risk factor, in Barrett’s we have previously shown gland epithelial phenotype diversity is increased prior to cancer progression^52^ and others have shown that genomic diversity follows a similar path^10,58^. We hypothesize that in this dysregulated microenvironment, the resulting impaired immune surveillance allows for the propagation of deleterious genomic aberrations (such as Chr18 amplifications) and eventual development of dysplasia and cancer.

Strikingly, NK cells were extremely rare in non-progressor patients, almost to the point where they were exclusive to pre-progressor and progressor patients (**Figure 1D**). Such a stark signature constitutes a very promising basis for a meaningful clinical test for risk stratification at early stages of BE. NK cells appear to play an important role in the pre- and progressive NDBE immune landscape showing strong attraction to most other immune cells. They appear as regulatory NK cells (lacking CD3 and expressing CD56^hi^, P63, CD11b and Annexin A1)^59^. Regulatory NK cells are known to express non-classical MHC class I molecules such as HLA-A, -B, - and -E^60^, an increase in expression of these genes was confirmed in our RNAseq data set (**Figure S13A**). NK cells appear to encroach into most CNs suggesting a broad repertoire of function. The mode of action of NK cells on the immune response in NDBE is not known but published evidence suggest that they can produce TGFβ^61^ and IL-10^62^ immune suppressive cytokines – notably, we observe an increase in the TGFβ signature (**Figure 5F**) and in TGFβ, TGFβI, TGFβR1/2/3 gene expression in both pre-progressors and progressors (**Figure S13B).** Additionally, NK cells are known to compete for binding to MHC class II antigens, thus preventing immune activation^63^ and even showing the ability to kill CD4^+^ T cells^64^; however, such a killing activity would likely be dependent on the expression of perforin which we did not detect in our study. The exact nature of the role of NK cells in BE progression therefore emerges as an important area of future research and clinical applications.

The role of intraepithelial immune cells in BE has never been studied. Yet, they represent an important immune sensing population of cells that are known to maintain homeostasis in the intestine. They are common in the small intestine (1 IEL per 10 epithelial cells) and the majority are CD8^+^ T cells^65^, however IELs are rarer in NDBE (2-3 cells per 100 epithelial cells, **Figure 2D**) in all three cohorts studied. We also detected intraepithelial CD4^+^ T cells, NK cells and M1 macrophages, suggesting that the epithelial layer is an active immune site hitherto unappreciated. Intraepithelial CD8^+^ T cells and CD4^+^ T cells were found in similar numbers in non-progressive NDBE but significantly more CD8^+^ IELs compared to CD4^+^ IELS were observed in pre-progressor and progressor NDBE samples. The ratio of CD8:CD4 IELs is therefore a strong indicator of patients on the pathway to cancer progression. The biological effect of altered T cell IEL ratios is unknown, but IELs are known to act as tumor suppressor cells^66,67^ and their dysregulation may play an important role in susceptibility to progression.

We demonstrated a strong positive correlation with chr18 amplifications and a negative correlation with chr9 deletions in progressor NDBE. While genomic alterations are well characterized in Barrett’s^8–10^, their association with the immune system is not. We observed an association between an increase in NK cells and neutrophil density and chr18 amplifications, even if a chr9 deletion is present. M1 macrophages and plasma cells on the other hand did not show the same association, indeed appeared randomly distributed with chr18 and ch9 alterations. This suggests that the spread of genetic abnormalities occurs later than M1 macrophage/plasma cell mediated inflammation. NK cells and neutrophils likely accrue in the progressor NDBE mucosa and are either precursors to genomic damage or accumulate as a result of it. The precise timeline of immune: genomic evolution cannot be determined here, but our data suggests they are dependent on each other.

Overall, our data demonstrate that the stromal-immune response alters prior to the development of overt cancer and is likely to play an important part in this process. More investigation into the timing of inflammation and the spread of genetic abnormalities in Barrett’s esophagus is required.

## Author contributions

Concept and funding: SM, GN, LF, SH, PG, TT

Lab based experiments: ML, JH, AP, MC, YD, SD

Computational analysis: ML, JH, YD

Pathology analysis: JC, MJ, MRJ, MN, MS, NAW, CMS

Manuscript preparation: All authors

Tissue collection and TMA preparation: JC, MRJ, MN, LF, RH, JE, VS, TJU, MM, CMS

## Supplemental figure Legends

**Figure S1. Representative CODEX images of tissue microarrays**. (**A**) Fluorescent imaging of 5 antibodies representing subpopulations of immune cells (CD8, red; CD4, green; CD20, yellow; HLADR, cyan; CD68, magenta). (**B**) Fluorescent imaging of 5 antibodies representing subpopulations of epithelial cells (MUC5AC, red; MUC2, green; MUC6, yellow; Defensin 5, cyan; PGA3, magenta). Tonsil controls are included in the bottom right-hand corner of each.

**Figure S2. CODEX expression dot plots**. Mean expression of CODEX markers (colour scale) and the fraction of cells in each marker (dot size) for (**A**) Non-progressor samples and (**B**) Pre-progressor samples.

**Figure S3. Cell maps for all identified immune cells**. (**A-C**) Non-progressor NDBE core maps coloured for each cell type. (**D**) Pre-progressor NDBE cores and (**E**) Progressor NDBE cores.

**Figure S4 Plasma cell characterisation using CODEX**. (**A**) Fraction of cells in non-progressor, pre-progressor and progressor metaplasia that are CD38^hi^ CD138^+^, CD38^lo^ CD138^+^, CD38^hi^ CD138^-^ or CD38^lo^ CD138^-^. (**B**) Representative examples of CODEX stains for CD38^hi^ and CD138^+^ with associated cell maps in non-progressor NDBE. (**C**) Pre-progressor NDBE, (**D**) Progressor NDBE. (*P<0.05, **P<0.01, ***P<0.001, ****P<0.0001, Wilcoxon rank sum test).

**Figure S5 Breakdown of CODEX cellular MMP9 expression by in non-progressor, pre-progressor and progressor metaplasia**. (**A**) Quantification of Lamina propria neutrophils between non-progressor, pre-progressor and progressor NDBE. (**B**) Representative CODEX staining and associated maps for MMP^hi^ and MMP9^lo^ neutrophils in non-progressor NDBE, (**C**) pre-progressor NDBE and (**D**) progressor NDBE. No other cell type expressed MMP9. (***P<0.001, ****P<0.0001, Wilcoxon rank sum test).

**Figure S6 Inferring cell identify in non-progressor, pre-progressor and progressor metaplasia from LCM RNAseq data**. (**A**) Expression of antibody mRNA expression across all cohorts. (**B**) Enrichr (<u>https://maayanlab.cloud/Enrichr/</u>) analysis revealed antibody mRNA was produced by B cells/plasma cells.(**C**) Complement gene expression is significantly across pre-progressor and progressor NBDE. (**D**) Enrichr analysis confirmed expression of complement genes by macrophages. (**E**) Genes known to be expressed by immunoregulatory NK cells show significant increase in expression in pre-progressor and progressor NBDE. (**F**) Enrichr analysis confirmed expression by NK cells. (**G**) S100A8-11 gene expression is also significantly increased in pre-progressor and progressor NDBE compared with non-progressors). (**H**) Enrichr analysis confirmed neutrophil expression of S100A8-11. (**I**) Expression of known dendritic cell markers representing myeloid and pre-dendritic cell populations. ITGAX could only be categorized as a general dendritic cell marker.(**J**) and (**K**) revealed that Enrichr analysis confirmed DC expression.). (*P<0.05, **P<0.01, ***P<0.001, ****P<0.0001, Wilcoxon rank sum test).

**Figure S7 Representative images of pairwise cell location and Ripley K function (transformed to L value) in non-progressor, pre-progressor and progressor metaplasia**. (**A**) Plasma cells: CD4+ Treg cells, (**B**) Plasma cells: CD8+ T cells and (**C**) plasma cells: CD4+ T cells. The number of cores showing attraction, repulsion, or neutrality used to calculate the interaction score is shown in the right panel.

**Figure S8. Cell interaction plots for other distances between cells**. (**Ai, Bi, Ci**) cell to cell interaction scores of all immune cells at 50 distance units for NP vs PP, PP vs P and NP vs P respectively. (**Aii, Bii, Cii**) interaction scores covering the entire cores and all distances were plotted. (**Aiii-v, Biii-v,Ciii-v**) Interaction score differences between cohorts were shown in a histogram(*=P<0.05, Fisher’s exact test).

**Figure S9.** Shannon diversity index of cell types across all neighborhoods in non-progressor, pre-progressor, and progressor NDBE.

**Figure S10**. Cell neighbourhood maps shown on all non-progressor (A-C), pre-progressor (D) and progressor (E) NDBE cores.

**Figure S11**. (A) Characterisation of CD38^hi^ CD138^+^ plasma cells across all cell neighbourhoods. (B) Characterisation of CD38^lo^ CD138^+^ plasmas cells across all cell neighbourhoods. Each compared between all cohorts. (*P<0.05, **P<0.01, ***P<0.001, ****P<0.0001, Wilcoxon rank sum test)

**Figure S12 Representative shallow WGS plots and P53 CODEX stains**. (**A**) Diploid plot. (**B**) Chr18 amplification plot. (**C**) Chr9 loss and Chr18 amplification plot. (**D**) Chr9 loss and Chr20 amplification plot. All CNVs that were not Chr9 loss or Chr18 amplifications were binned as ‘CNVs’ in **Figure 7. (E-H)** representative p53 staining with H&Es on the same section on samples that were either p53 negative or positive.

**Figure S13. Gene expression for non-classical HLAs and TGF*β*family mRNA expression**. The log2 mean counts were compared with NP, PP & P samples for (A) HLA-A, HLA-C and HLA-E mRNA and (B) TGFB1, TGFBI, TGBR1, TGFBR2 & TGFBR3. (***=P<0.001, **=P<0.01, * P<0.05, Wilcoxon rank sum test).

## References

1. Giroux, V., and Rustgi, A.K. (2017). Metaplasia: tissue injury adaptation and a precursor to the dysplasia-cancer sequence. Nat Rev Cancer 17, 594–604. 10.1038/nrc.2017.68.

2. Colotta, F., Allavena, P., Sica, A., Garlanda, C., and Mantovani, A. (2009). Cancer-related inflammation, the seventh hallmark of cancer: links to genetic instability. Carcinogenesis 30, 1073–1081. 10.1093/carcin/bgp127.

3. Galassi, C., Chan, T.A., Vitale, I., and Galluzzi, L. (2024). The hallmarks of cancer immune evasion. Cancer Cell 42, 1825–1863. 10.1016/j.ccell.2024.09.010.

4. Curtius, K., Rubenstein, J.H., Chak, A., and Inadomi, J.M. (2020). Computational modelling suggests that Barrett’s oesophagus may be the precursor of all oesophageal adenocarcinomas. Gut 70, 1435–1440. 10.1136/gutjnl-2020-321598.

5. McDonald, S.A.C., Graham, T.A., Lavery, D., Wright, N.A., and Jansen, M. (2015). The Barretts gland in phenotype space. Cellular and Molecular Gastroenterology and Hepatology 1, 41–54.

6. Dulak, A.M., Stojanov, P., Peng, S., Lawrence, M.S., Fox, C., Stewart, C., Bandla, S., Imamura, Y., Schumacher, S.E., Shefler, E., et al. (2013). Exome and whole-genome sequencing of esophageal adenocarcinoma identifies recurrent driver events and mutational complexity. Nat Genet 45, 478–486. 10.1038/ng.2591.

7. Ross-Innes, C.S., Becq, J., Warren, A., Cheetham, R.K., Northen, H., O’Donovan, M., Malhotra, S., di Pietro, M., Ivakhno, S., He, M., et al. (2015). Whole-genome sequencing provides new insights into the clonal architecture of Barrett’s esophagus and esophageal adenocarcinoma. Nat Genet 47, 1038–1046.

8. Paulson, T.G., Galipeau, P.C., Oman, K.M., Sanchez, C.A., Kuhner, M.K., Smith, L.P., Hadi, K., Shah, M., Arora, K., Shelton, J., et al. (2022). Somatic whole genome dynamics of precancer in Barrett’s esophagus reveals features associated with disease progression. Nat Commun 13, 2300.

9. Killcoyne, S., Gregson, E., Wedge, D.C., Woodcock, D.J., Eldridge, M.D., de la Rue, R., Miremadi, A., Abbas, S., Blasko, A., Kosmidou, C., et al. (2020). Genomic copy number predicts esophageal cancer years before transformation. Nat Med 26, 1726–1732. 10.1038/s41591-020-1033-y.

10. Maley, C.C., Galipeau, P.C., Finley, J.C., Wongsurawat, V.J., Li, X., Sanchez, C.A., Paulson, T.G., Blount, P.L., Risques, R.-A., Rabinovitch, P.S., and Reid, B.J. (2006). Genetic clonal diversity predicts progression to esophageal adenocarcinoma. Nat Genet 38, 468–473. 10.1038/ng1768.

11. Killcoyne, S., and Fitzgerald, R.C. (2021). Evolution and progression of Barrett’s oesophagus to oesophageal cancer. Nat Rev Cancer 21, 731–741. 10.1038/s41568-021-00400-x.

12. Redston, M., Noffsinger, A., Kim, A., Akarca, F.G., Rara, M., Stapleton, D., Nowden, L., Lash, R., Bass, A.J., and Stachler, M.D. (2022). Abnormal TP53 Predicts Risk of Progression in Patients With Barrett’s Esophagus Regardless of a Diagnosis of Dysplasia. Gastroenterology 162, 468–481. 10.1053/j.gastro.2021.10.038.

13. Weaver, J.M., Ross-Innes, C.S., Shannon, N., Lynch, A.G., Forshew, T., Barbera, M., Murtaza, M., Ong, C.A., Lao-Sirieix, P., Dunning, M.J., et al. (2014). Ordering of mutations in preinvasive disease stages of esophageal carcinogenesis. Nat Genet 46, 837–843. 10.1038/ng.3013.

14. Timmer, M.R., Martinez, P., Lau, C.T., Westra, W.M., Calpe, S., Rygiel, A.M., Rosmolen, W.D., Meijer, S.L., Ten Kate, F.J., Dijkgraaf, M.G., et al. (2015). Derivation of genetic biomarkers for cancer risk stratification in Barrett’s oesophagus: a prospective cohort study. Gut. 10.1136/gutjnl-2015-309642.

15. Tambunting, L., Kelleher, D., and Duggan, S.P. (2022). The Immune Underpinnings of Barrett’s-Associated Adenocarcinogenesis: a Retrial of Nefarious Immunologic Co-Conspirators. Cell Mol Gastroenterol Hepatol 13, 1297–1315. 10.1016/j.jcmgh.2022.01.023.

16. Power, R., Lowery, M.A., Reynolds, J.V., and Dunne, M.R. (2020). The Cancer-Immune Set Point in Oesophageal Cancer. Front Oncol 10, 891. 10.3389/fonc.2020.00891.

17. Fitzgerald, R.C., Onwuegbusi, B.A., Bajaj-Elliott, M., Saeed, I.T., Burnham, W.R., and Farthing, M.J. (2002). Diversity in the oesophageal phenotypic response to gastro-oesophageal reflux: immunological determinants. Gut 50, 451–459. 10.1136/gut.50.4.451.

18. van Sandick, J.W., Boermeester, M.A., Gisbertz, S.S., ten Berge, I.J., Out, T.A., van der Pouw Kraan, T.C., and van Lanschot, J.J. (2003). Lymphocyte subsets and T(h)1/T(h)2 immune responses in patients with adenocarcinoma of the oesophagus or oesophagogastric junction: relation to pTNM stage and clinical outcome. Cancer Immunol Immunother 52, 617–624. 10.1007/s00262-003-0406-7.

19. Sundaram, S., Kim, E.N., Jones, G.M., Sivagnanam, S., Tripathi, M., Miremadi, A., Di Pietro, M., Coussens, L.M., Fitzgerald, R.C., Chang, Y.H., and Zhuang, L. (2022). Deciphering the Immune Complexity in Esophageal Adenocarcinoma and Pre-Cancerous Lesions With Sequential Multiplex Immunohistochemistry and Sparse Subspace Clustering Approach. Front Immunol 13, 874255. 10.3389/fimmu.2022.874255.

20. Lagisetty, K.H., McEwen, D.P., Nancarrow, D.J., Schiebel, J.G., Ferrer-Torres, D., Ray, D., Frankel, T.L., Lin, J., Chang, A.C., Kresty, L.A., and Beer, D.G. (2021). Immune determinants of Barrett’s progression to esophageal adenocarcinoma. JCI Insight 6. 10.1172/jci.insight.143888.

21. Strasser, M.K., Gibbs, D.L., Gascard, P., Bons, J., Hickey, J.W., Schurch, C.M., Tan, Y., Black, S., Chu, P., Ozkan, A., et al. (2023). Concerted epithelial and stromal changes during progression of Barrett’s Esophagus to invasive adenocarcinoma exposed by multi-scale, multi-omics analysis. bioRxiv. 10.1101/2023.06.08.544265.

22. Li, X., Galipeau, P.C., Paulson, T.G., Sanchez, C.A., Arnaudo, J., Liu, K., Sather, C.L., Kostadinov, R.L., Odze, R.D., Kuhner, M.K., et al. (2014). Temporal and spatial evolution of somatic chromosomal alterations: a case-cohort study of Barrett’s esophagus. Cancer prevention research 7, 114–127.

23. Black, S., Phillips, D., Hickey, J.W., Kennedy-Darling, J., Venkataraaman, V.G., Samusik, N., Goltsev, Y., Schurch, C.M., and Nolan, G.P. (2021). CODEX multiplexed tissue imaging with DNA-conjugated antibodies. Nat Protoc 16, 3802–3835. 10.1038/s41596-021-00556-8.

24. Lee, M.Y., Bedia, J.S., Bhate, S.S., Barlow, G.L., Phillips, D., Fantl, W.J., Nolan, G.P., and Schurch, C.M. (2022). CellSeg: a robust, pre-trained nucleus segmentation and pixel quantification software for highly multiplexed fluorescence images. BMC Bioinformatics 23, 46. 10.1186/s12859-022-04570-9.

25. Hickey, J.W., Tan, Y., Nolan, G.P., and Goltsev, Y. (2021). Strategies for Accurate Cell Type Identification in CODEX Multiplexed Imaging Data. Front Immunol 12, 727626. 10.3389/fimmu.2021.727626.

26. Samusik, N., Good, Z., Spitzer, M.H., Davis, K.L., and Nolan, G.P. (2016). Automated mapping of phenotype space with single-cell data. Nat Methods 13, 493–496. 10.1038/nmeth.3863.

27. Amezquita, R.A., Lun, A.T.L., Becht, E., Carey, V.J., Carpp, L.N., Geistlinger, L., Marini, F., Rue-Albrecht, K., Risso, D., Soneson, C., et al. (2020). Publisher Correction: Orchestrating single-cell analysis with Bioconductor. Nat Methods 17, 242. 10.1038/s41592-019-0700-8.

28. Haghverdi, L., Lun, A.T.L., Morgan, M.D., and Marioni, J.C. (2018). Batch effects in single-cell RNA-sequencing data are corrected by matching mutual nearest neighbors. Nat Biotechnol 36, 421–427. 10.1038/nbt.4091.

29. McCarthy, D.J., Campbell, K.R., Lun, A.T., and Wills, Q.F. (2017). Scater: pre-processing, quality control, normalization and visualization of single-cell RNA-seq data in R. Bioinformatics 33, 1179–1186. 10.1093/bioinformatics/btw777.

30. Wickham, R. (2016). Elegant Graphics for Data Analysis (Springer-Verlag New York).

31. Kassambara, A. (2023). rstatix: Pipe-Friendly Framework for Basic Statistical Tests. https://rpkgs.datanovia.com/rstatix/.

32. Schurch, C.M., Bhate, S.S., Barlow, G.L., Phillips, D.J., Noti, L., Zlobec, I., Chu, P., Black, S., Demeter, J., McIlwain, D.R., et al. (2020). Coordinated Cellular Neighborhoods Orchestrate Antitumoral Immunity at the Colorectal Cancer Invasive Front. Cell 182, 1341–1359.

33. Bhate, S.S., Barlow, G.L., Schurch, C.M., and Nolan, G.P. (2022). Tissue schematics map the specialization of immune tissue motifs and their appropriation by tumors. Cell Syst 13, 109–130 e106. 10.1016/j.cels.2021.09.012.

34. Baddeley, A., Rubak, E., and Turner, R. (2015). Spatial Point Patterns: Methodology and Applications with R (Chapman and Hall/CRC Press, London).

35. Diggle, P.J., Mateu, J., and Clough, H.E. (2000). A Comparison between Parametric and Non-Parametric Approaches to the Analysis of Replicated Spatial Point Patterns. Advances in Applied Probability 32, 331–343.

36. Foley, J.W., Zhu, C., Jolivet, P., Zhu, S.X., Lu, P., Meaney, M.J., and West, R.B. (2019). Gene expression profiling of single cells from archival tissue with laser-capture microdissection and Smart-3SEQ. Genome Res 29, 1816–1825. 10.1101/gr.234807.118.

37. Andrews, S. (2010). FastQC: a quality control tool for high throughput sequence data. http://www.bioinformatics.babraham.ac.uk/projects/fastqc.

38. Bolger, A.M., Lohse, M., and Usadel, B. (2014). Trimmomatic: a flexible trimmer for Illumina sequence data. Bioinformatics 30, 2114–2120. 10.1093/bioinformatics/btu170.

39. Pebesma, E., and Bivand, R. (2023). Spatial Data Science: With applications in R (Chapman and Hall/CRC). 10.1201/9780429459016.

40. Liao, Y., Smyth, G.K., and Shi, W. (2019). The R package Rsubread is easier, faster, cheaper and better for alignment and quantification of RNA sequencing reads. Nucleic Acids Res 47, e47. 10.1093/nar/gkz114.

41. Krijthe, J.H. (2015). Rtsne: T-Distributed Stochastic Neighbor Embedding using Barnes-Hut Implementation. https://github.com/jkrijthe/Rtsne.

42. Bolar, K. (2019). STAT: Interactive Document for Working with Basic Statistical Analysis. https://CRAN.R-project.org/package=STAT.

43. Love, M.I., Huber, W., and Anders, S. (2014). Moderated estimation of fold change and dispersion for RNA-seq data with DESeq2. Genome Biol 15, 550. 10.1186/s13059-014-0550-8.

44. Hanzelmann, S., Castelo, R., and Guinney, J. (2013). GSVA: gene set variation analysis for microarray and RNA-seq data. BMC Bioinformatics 14, 7. 10.1186/1471-2105-14-7.

45. Langfelder, P., and Horvath, S. (2008). WGCNA: an R package for weighted correlation network analysis. BMC Bioinformatics 9, 559. 10.1186/1471-2105-9-559.

46. Li, H., and Durbin, R. (2009). Fast and accurate short read alignment with Burrows-Wheeler transform. Bioinformatics 25, 1754–1760. 10.1093/bioinformatics/btp324.

47. Van der Auwera, G. (2020). Genomics in the Cloud (O’Reilly Media Inc).

48. Scheinin, I., Sie, D., Bengtsson, H., van de Wiel, M.A., Olshen, A.B., van Thuijl, H.F., van Essen, H.F., Eijk, P.P., Rustenburg, F., Meijer, G.A., et al. (2014). DNA copy number analysis of fresh and formalin-fixed specimens by shallow whole-genome sequencing with identification and exclusion of problematic regions in the genome assembly. Genome Res 24, 2022–2032. 10.1101/gr.175141.114.

49. Poell, J.B., Mendeville, M., Sie, D., Brink, A., Brakenhoff, R.H., and Ylstra, B. (2019). ACE: absolute copy number estimation from low-coverage whole-genome sequencing data. Bioinformatics 35, 2847–2849. 10.1093/bioinformatics/bty1055.

50. Vonwirth, V., Bulbul, Y., Werner, A., Echchannaoui, H., Windschmitt, J., Habermeier, A., Ioannidis, S., Shin, N., Conradi, R., Bros, M., et al. (2020). Inhibition of Arginase 1 Liberates Potent T Cell Immunostimulatory Activity of Human Neutrophil Granulocytes. Front Immunol 11, 617699. 10.3389/fimmu.2020.617699.

51. Ardi, V.C., Kupriyanova, T.A., Deryugina, E.I., and Quigley, J.P. (2007). Human neutrophils uniquely release TIMP-free MMP-9 to provide a potent catalytic stimulator of angiogenesis. Proc Natl Acad Sci U S A 104, 20262–20267. 10.1073/pnas.0706438104.

52. Evans, J.A., Carlotti, E., Lin, M.L., Hackett, R.J., Haughey, M.J., Passman, A.M., Dunn, L., Elia, G., Porter, R.J., McLean, M.H., et al. (2022). Clonal Transitions and Phenotypic Evolution in Barrett’s Esophagus. Gastroenterology 162, 1197–1209 e1113.

53. Delfini, M., Stakenborg, N., Viola, M.F., and Boeckxstaens, G. (2022). Macrophages in the gut: Masters in multitasking. Immunity 55, 1530–1548. 10.1016/j.immuni.2022.08.005.

54. Bagaev, A., Kotlov, N., Nomie, K., Svekolkin, V., Gafurov, A., Isaeva, O., Osokin, N., Kozlov, I., Frenkel, F., Gancharova, O., et al. (2021). Conserved pan-cancer microenvironment subtypes predict response to immunotherapy. Cancer Cell 39, 845–865 e847. 10.1016/j.ccell.2021.04.014.

55. Leedham, S.J., Preston, S.L., McDonald, S.A.C., Elia, G., Bhandari, P., Poller, D., Harrison, R., Novelli, M.R., Jankowski, J.A., and Wright, N.A. (2008). Individual crypt genetic heterogeneity and the origin of metaplastic glandular epithelium in human Barrett’s oesophagus. Gut 57, 1041–1048.

56. Paulson, T.G., Galipeau, P.C., Xu, L., Kissel, H.D., Li, X., Blount, P.L., Sanchez, C.A., Odze, R.D., and Reid, B.J. (2008). p16 mutation spectrum in the premalignant condition Barrett’s esophagus. PLoS ONE 3, e3809. 10.1371/journal.pone.0003809.

57. Ganguli, P., Basanta, C.C., Armero, A., Zahra, A., Sagrado, M., Devonshire, G., Freeman, A., Spencer, J., Fitzgerald, R.C., and Ciccarelli, F.D. (2024). Context-dependent effects of CDK2NA and other 9p21 gene losses during the evolution of oesophageal cancer. bioRxiv. 10.1101/2024.01.24.576991.

58. Martinez, P., Timmer, M.R., Lau, C.T., Calpe, S., Sancho-Serra Mdel, C., Straub, D., Baker, A.M., Meijer, S.L., Kate, F.J., Mallant-Hent, R.C., et al. (2016). Dynamic clonal equilibrium and predetermined cancer risk in Barrett’s oesophagus. Nat Commun 7, 12158.

59. Jiang, H., and Jiang, J. (2023). Balancing act: the complex role of NK cells in immune regulation. Front Immunol 14, 1275028. 10.3389/fimmu.2023.1275028.

60. Pegram, H.J., Andrews, D.M., Smyth, M.J., Darcy, P.K., and Kershaw, M.H. (2011). Activating and inhibitory receptors of natural killer cells. Immunol Cell Biol 89, 216–224. 10.1038/icb.2010.78.

61. Gray, J.D., Hirokawa, M., Ohtsuka, K., and Horwitz, D.A. (1998). Generation of an inhibitory circuit involving CD8+ T cells, IL-2, and NK cell-derived TGF-beta: contrasting effects of anti-CD2 and anti-CD3. J Immunol 160, 2248–2254.

62. Mehrotra, P.T., Donnelly, R.P., Wong, S., Kanegane, H., Geremew, A., Mostowski, H.S., Furuke, K., Siegel, J.P., and Bloom, E.T. (1998). Production of IL-10 by human natural killer cells stimulated with IL-2 and/or IL-12. J Immunol 160, 2637–2644.

63. Nakayama, M., Takeda, K., Kawano, M., Takai, T., Ishii, N., and Ogasawara, K. (2011). Natural killer (NK)-dendritic cell interactions generate MHC class II-dressed NK cells that regulate CD4+ T cells. Proc Natl Acad Sci U S A 108, 18360–18365. 10.1073/pnas.1110584108.

64. Fort, M.M., Leach, M.W., and Rennick, D.M. (1998). A role for NK cells as regulators of CD4+ T cells in a transfer model of colitis. J Immunol 161, 3256–3261.

65. Olivares-Villagomez, D., and Olivares-Van Kaer, L., (2018). Intestinal Intraepithelial Lymphocytes: Sentinels of the Mucosal Barrier. Trends Immunol 39, 264–275. 10.1016/j.it.2017.11.003.

66. Huang, J., Zhang, X., Xu, H., Fu, L., Liu, Y., Zhao, J., Huang, J., Song, Z., Zhu, M., Fu, Y.X., et al. (2024). Intraepithelial lymphocytes promote intestinal regeneration through CD160/HVEM signaling. Mucosal Immunol 17, 257–271. 10.1016/j.mucimm.2024.02.004.

67. Roberts, A.I., O’Connell, S.M., and Ebert, E.C. (1993). Intestinal intraepithelial lymphocytes bind to colon cancer cells by HML-1 and CD11a. Cancer Res 53, 1608–1611.

