## Supplementary figures and images for "Alterations of the composition and spatial organization of the microenvironment following non-dysplastic Barrett’s esophagus through progression to cancer"

### Supplemental figure 1

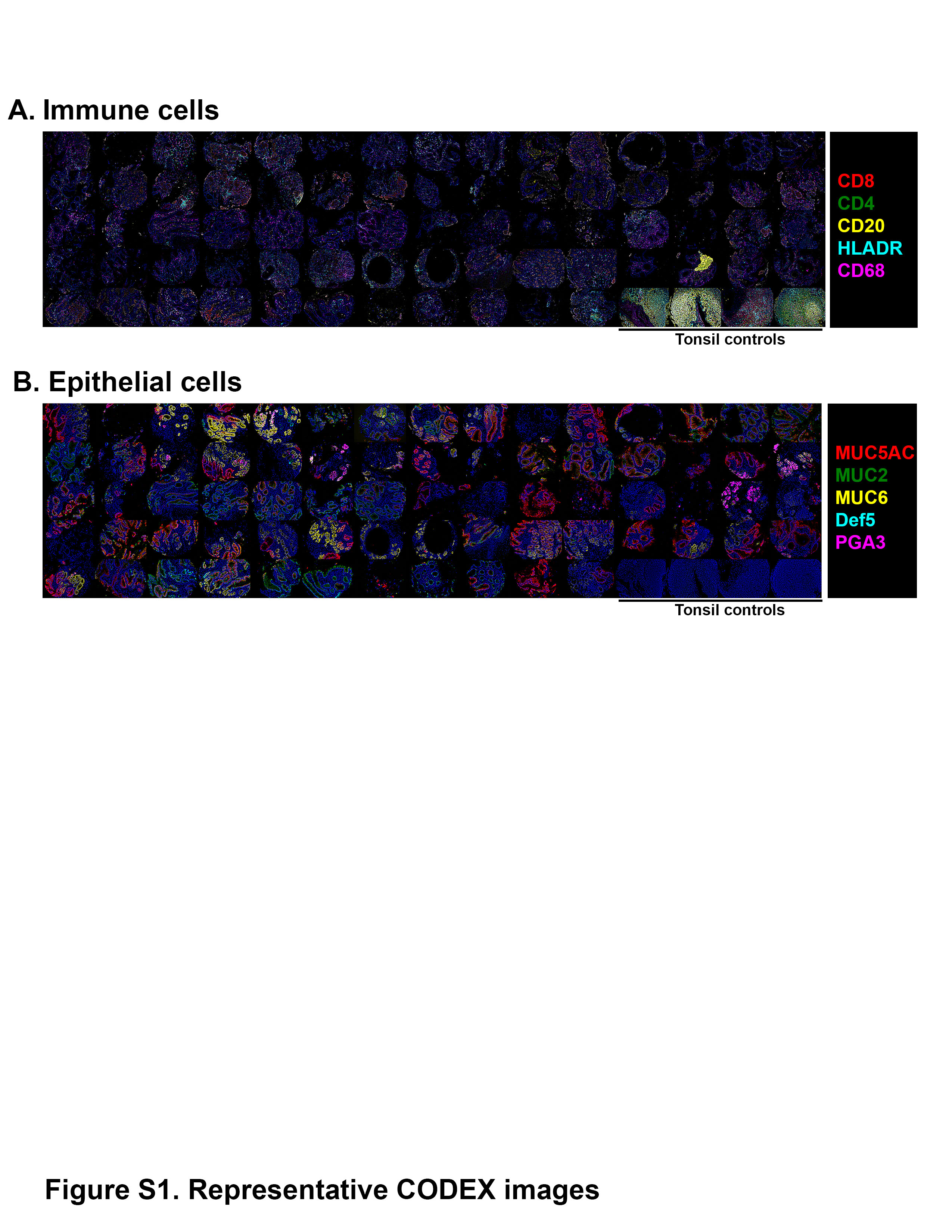

### Supplemental figure 2

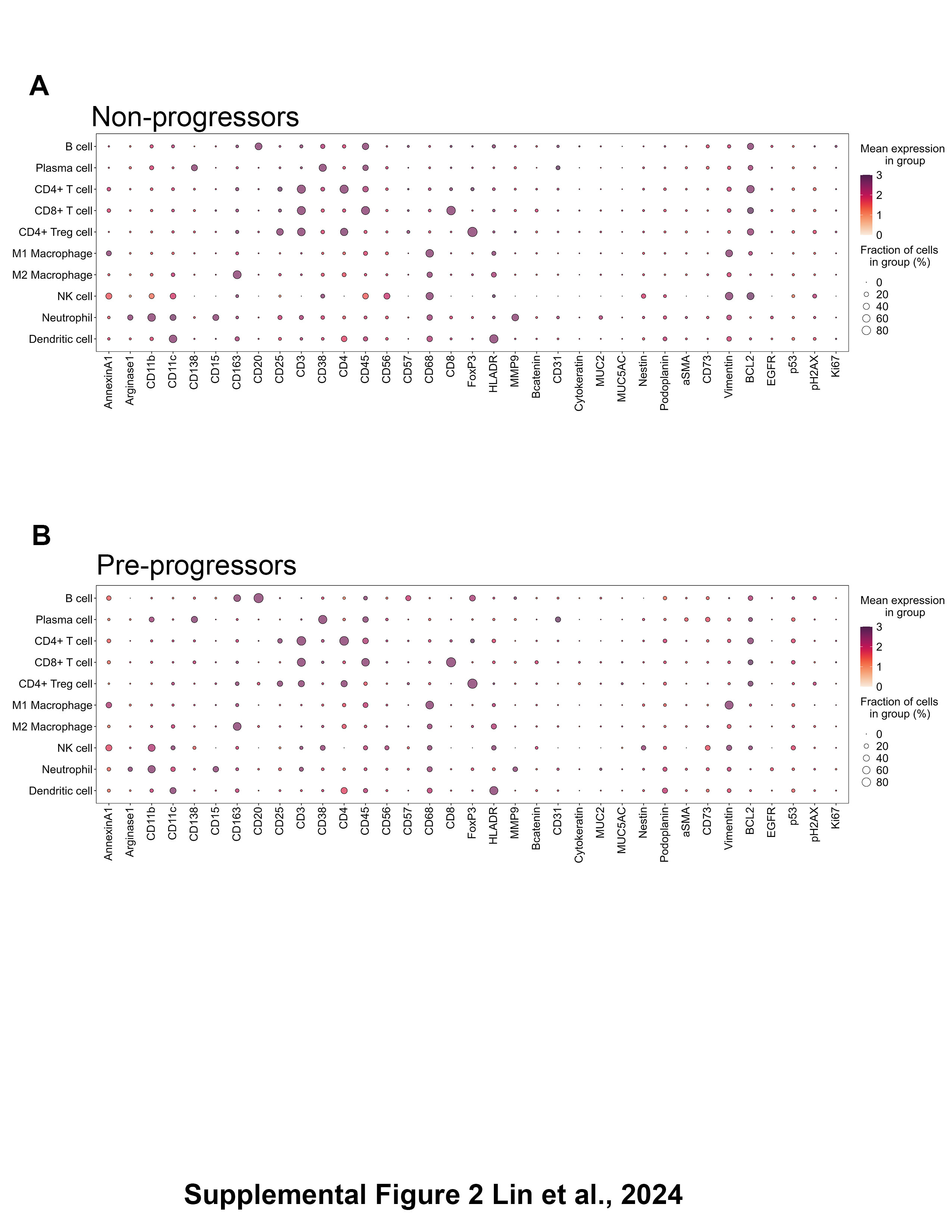

### Supplemental figure 3A

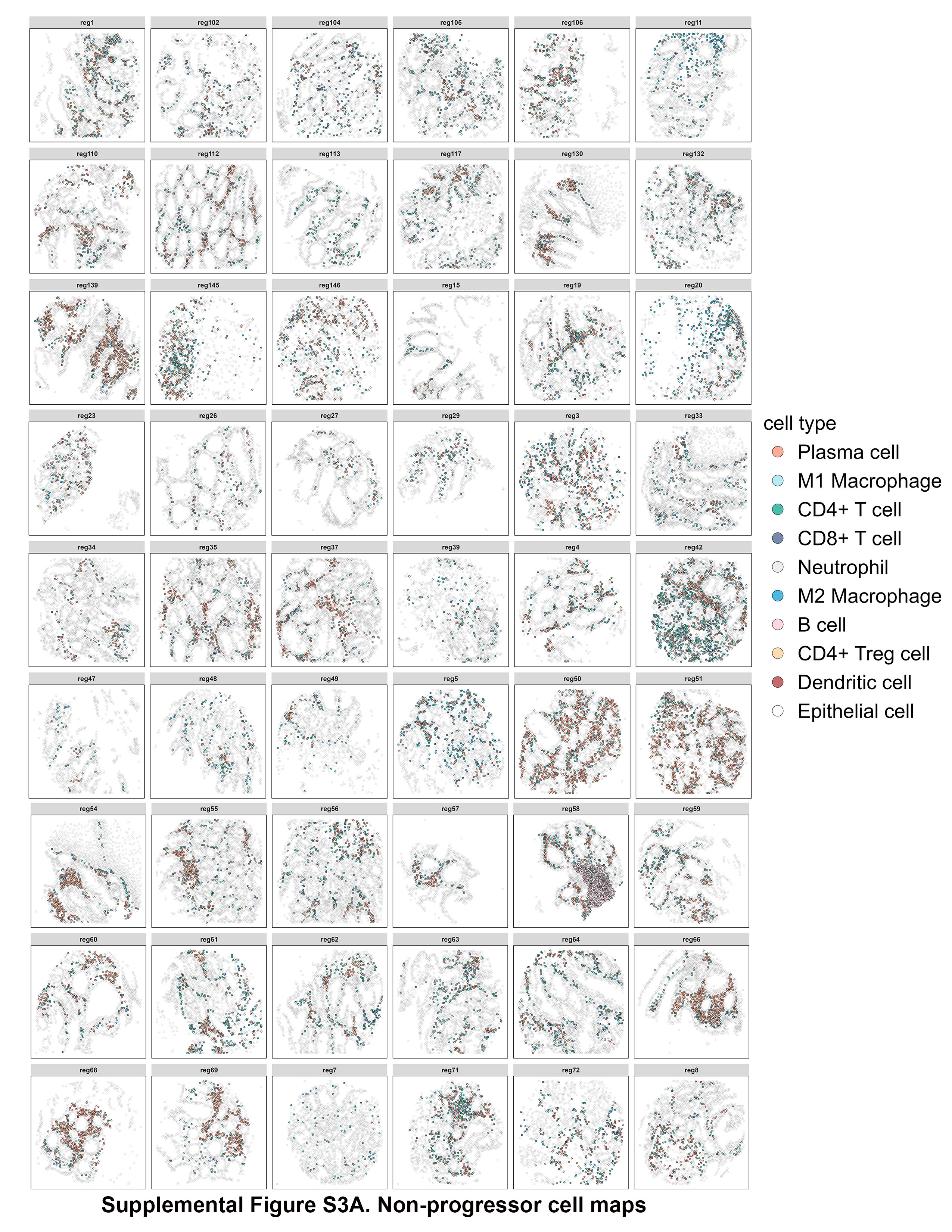

### Supplemental figure 3B

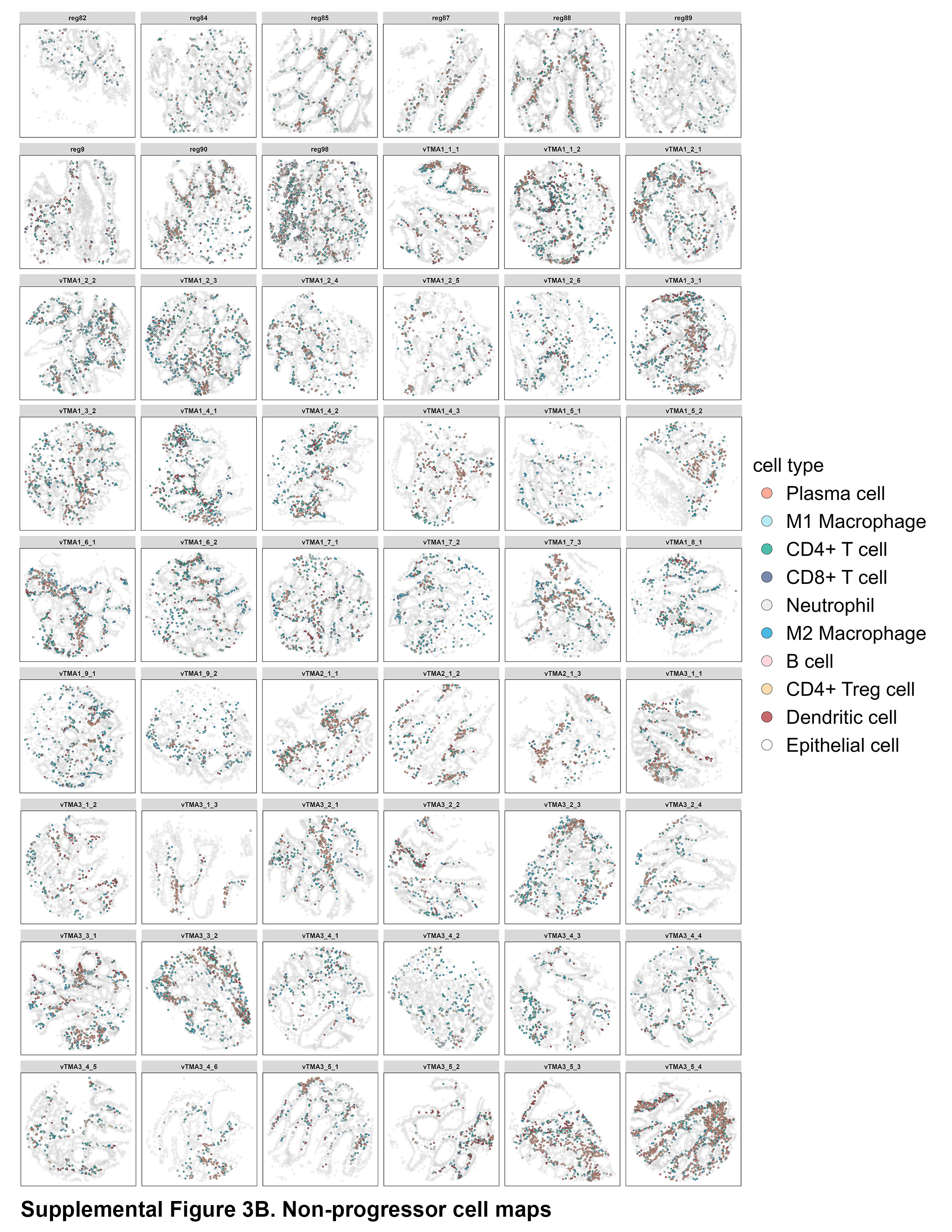

### Supplemental figure 3C

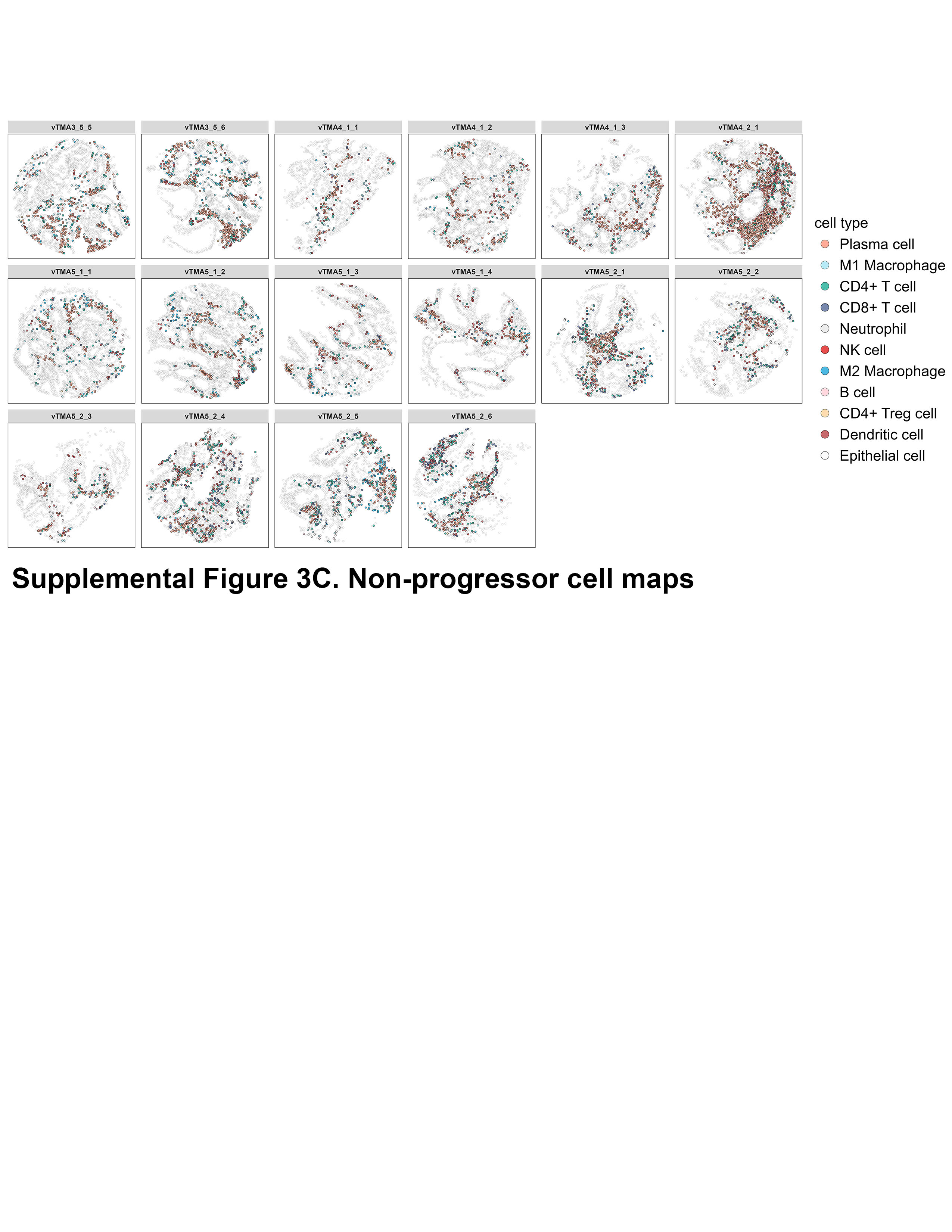

### Supplemental figure 3D

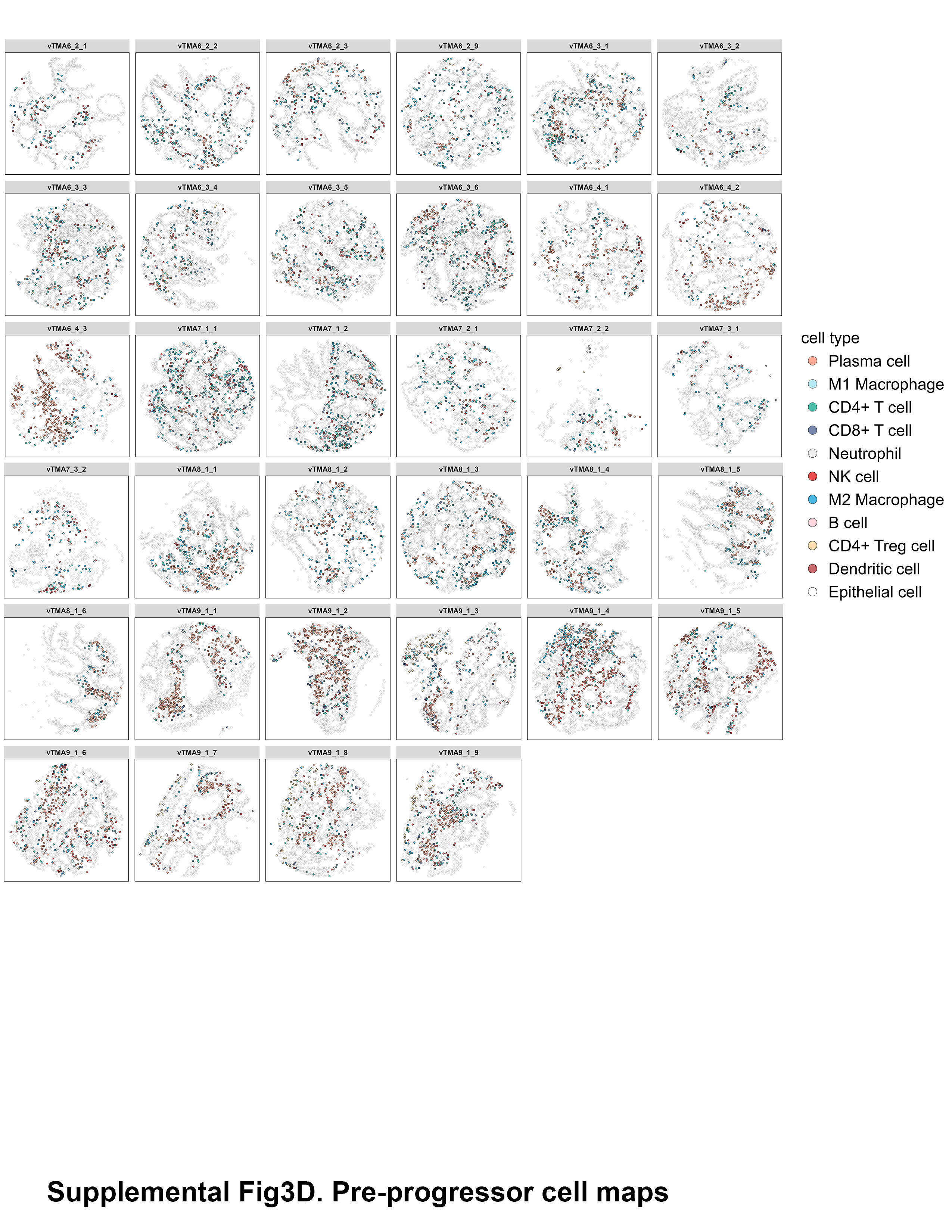

### Supplemental figure 3E

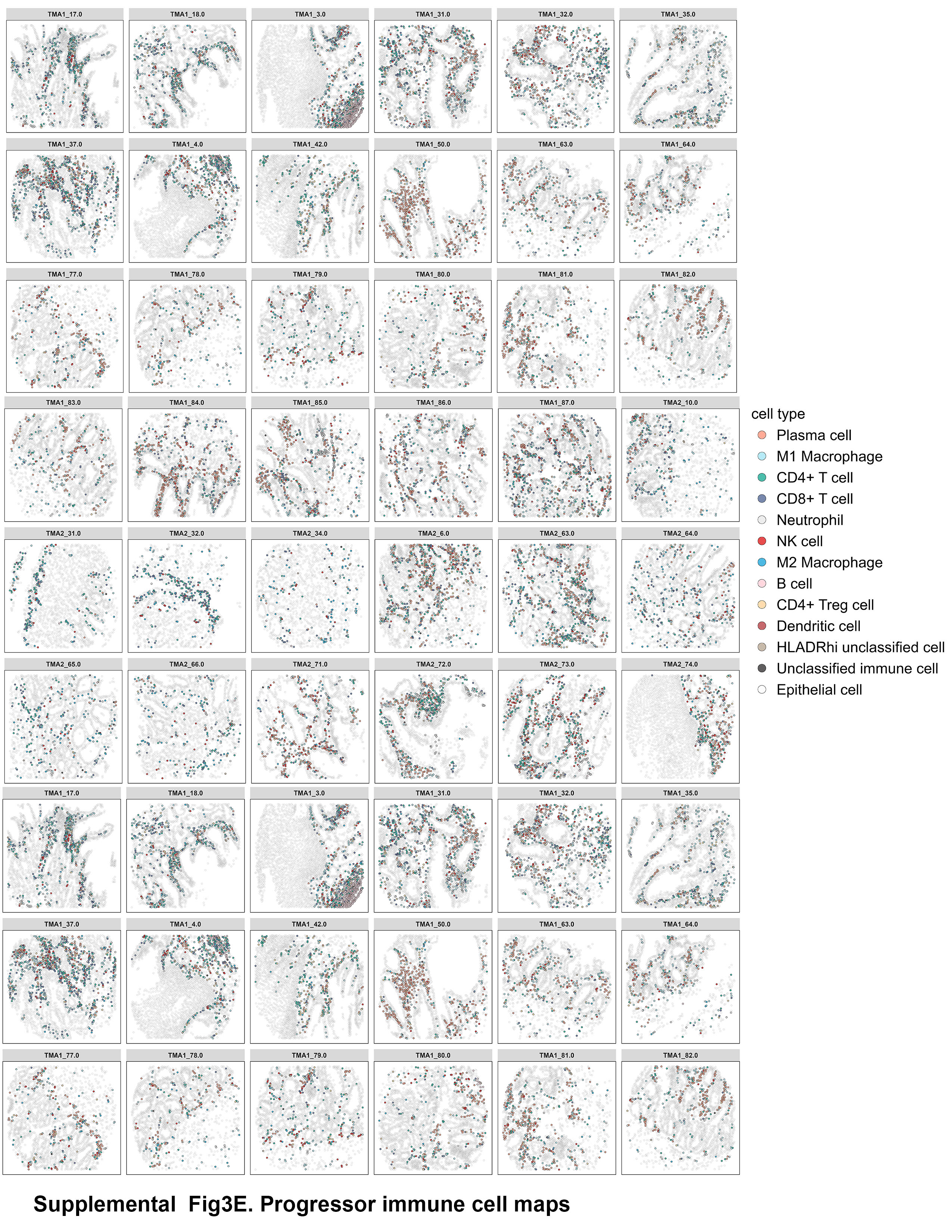

### Supplemental figure 4

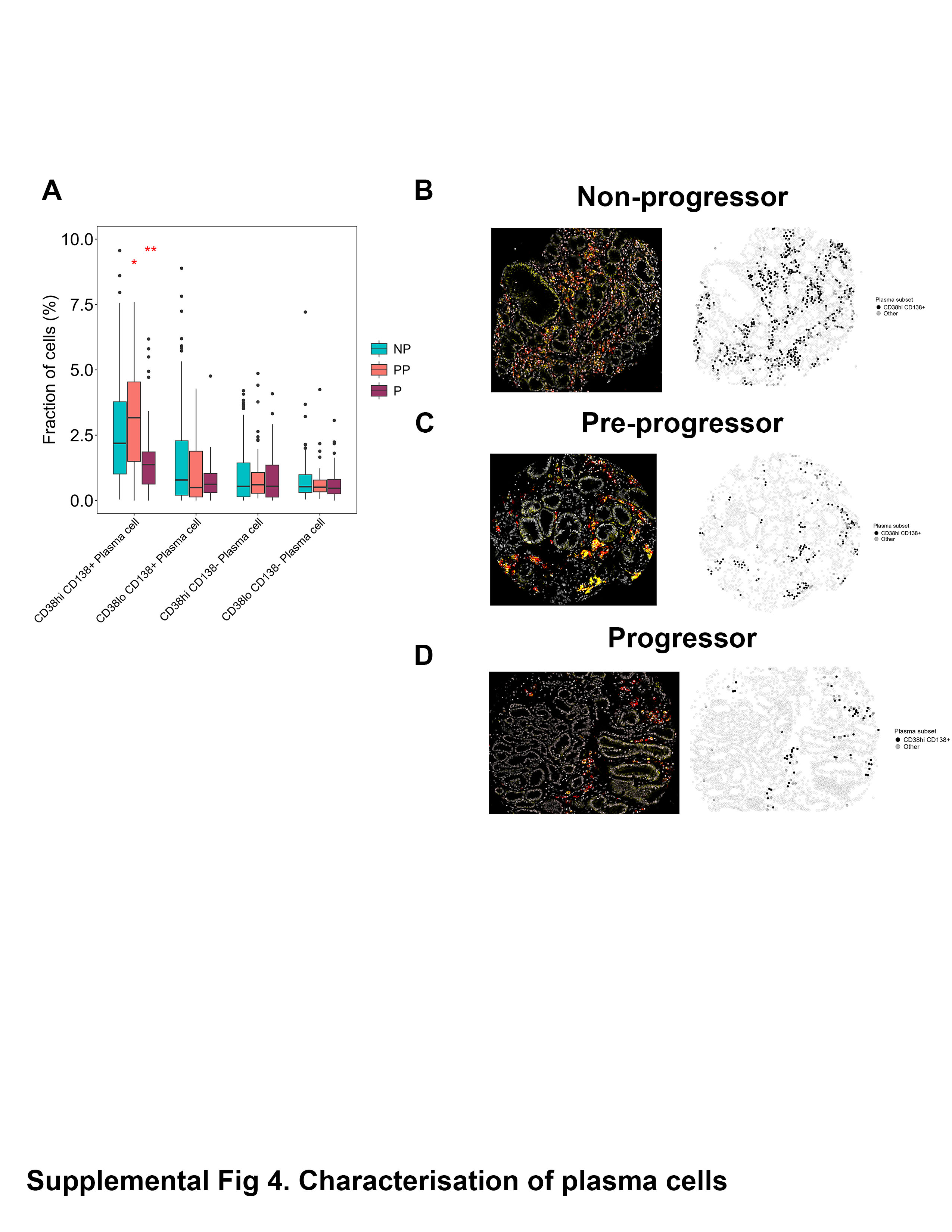

### Supplemental figure 5

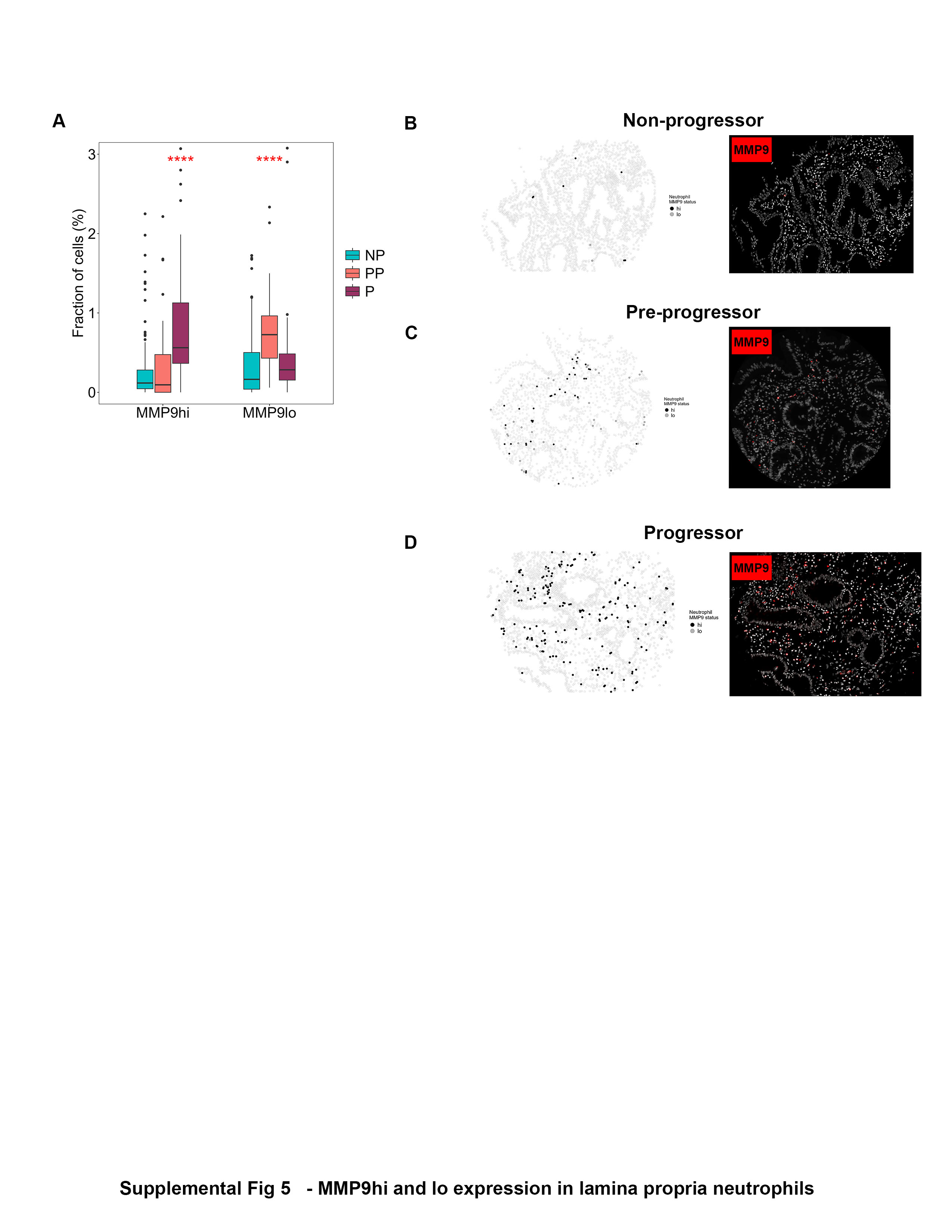

### Supplemental figure 6

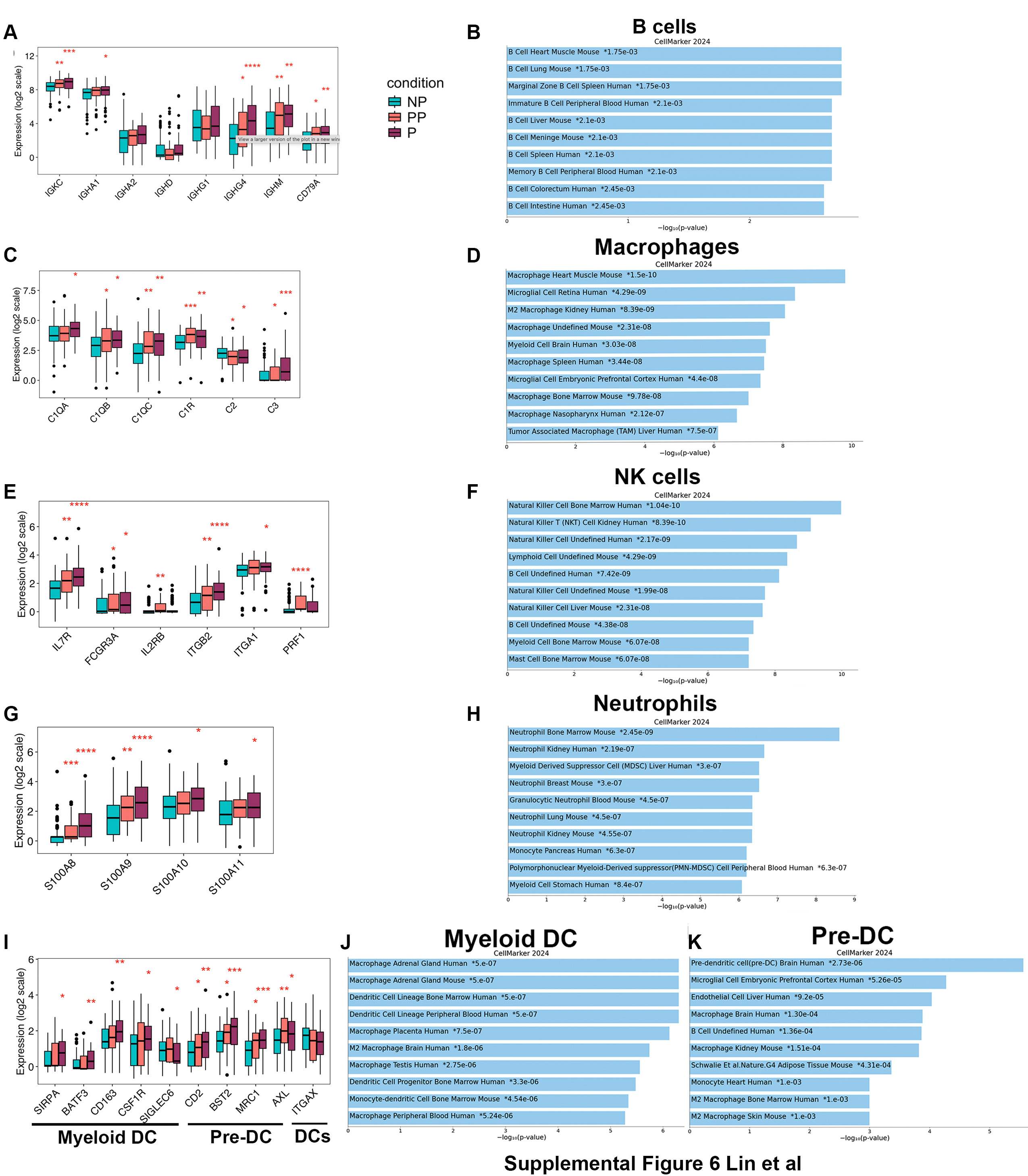

### Supplemental figure 7

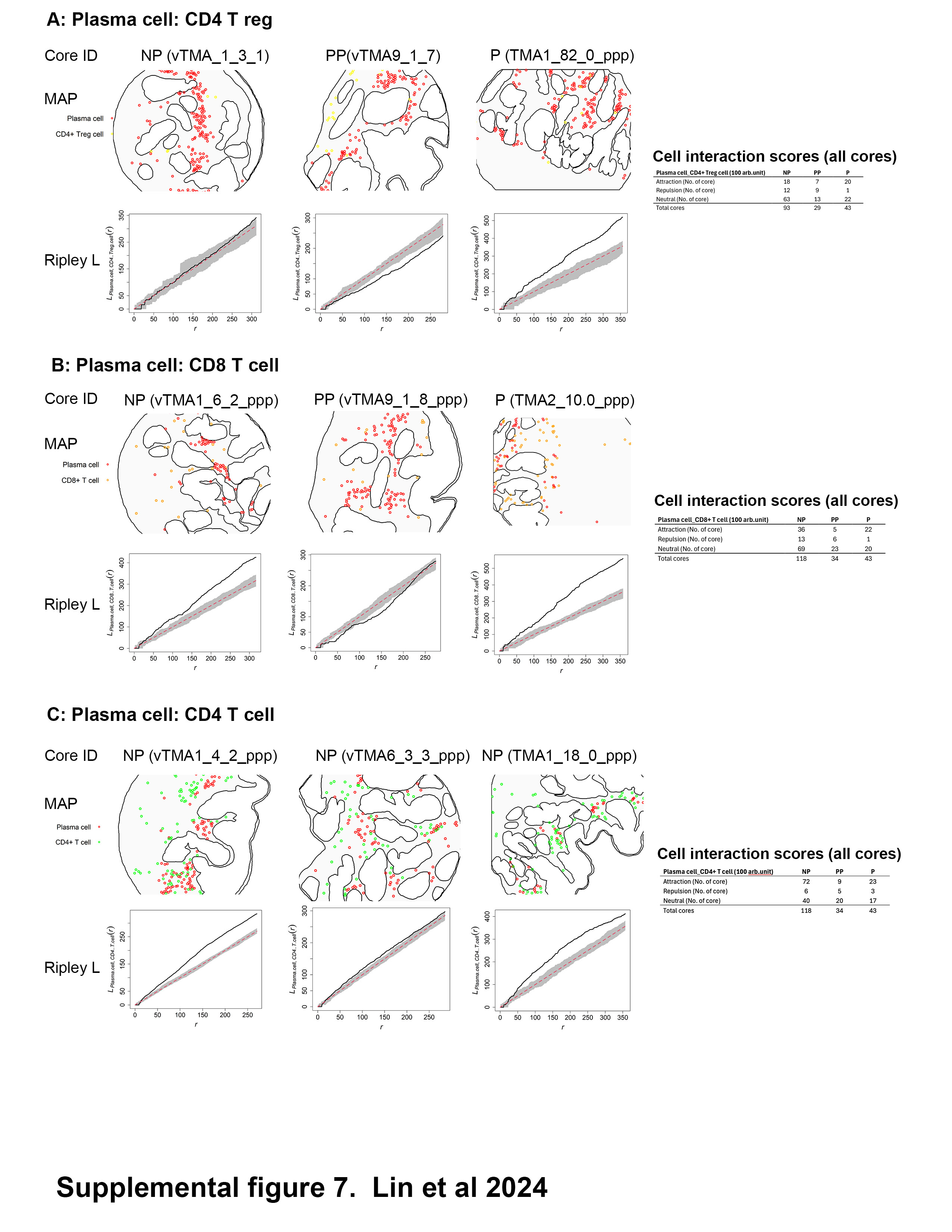

### Supplemental figure 8

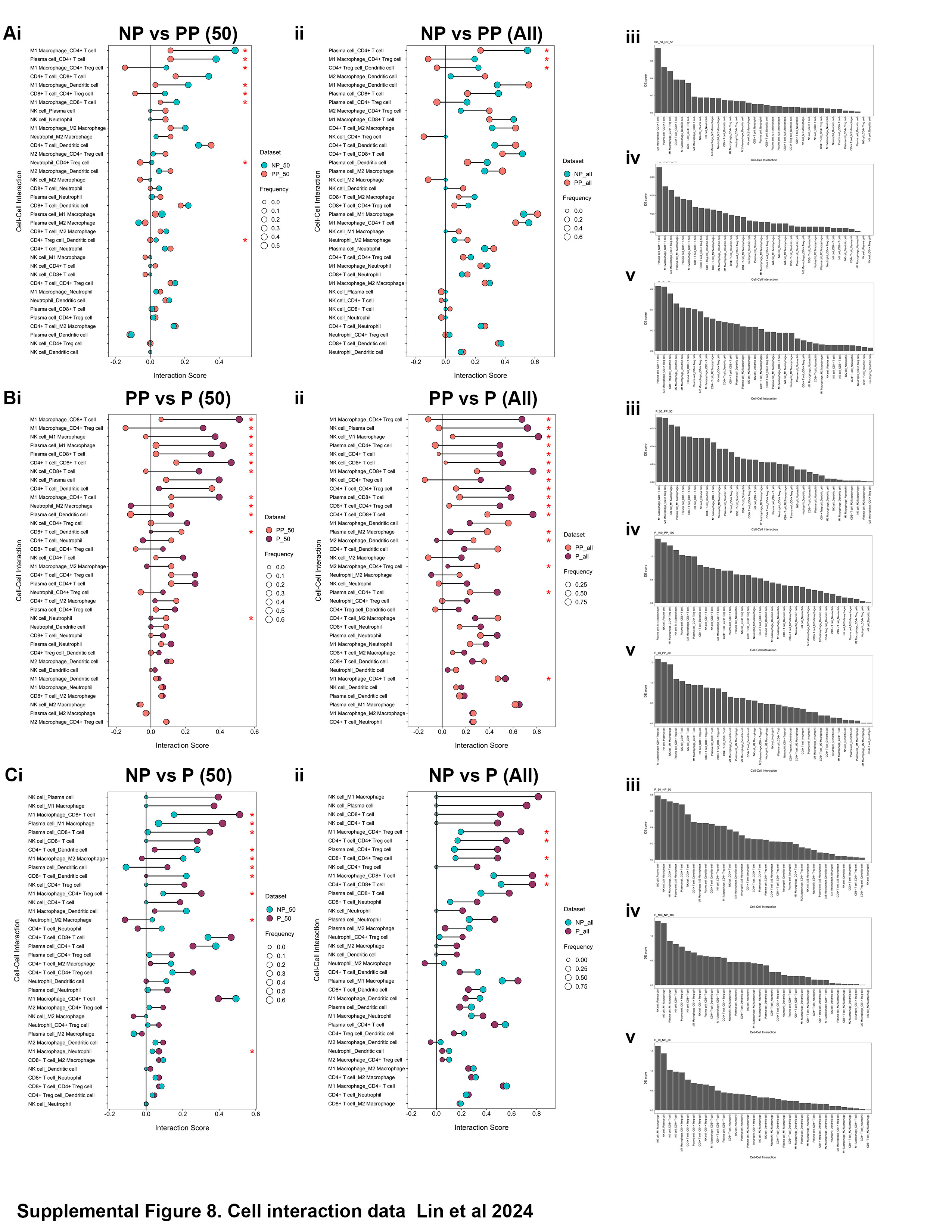

### Supplemental figure 9

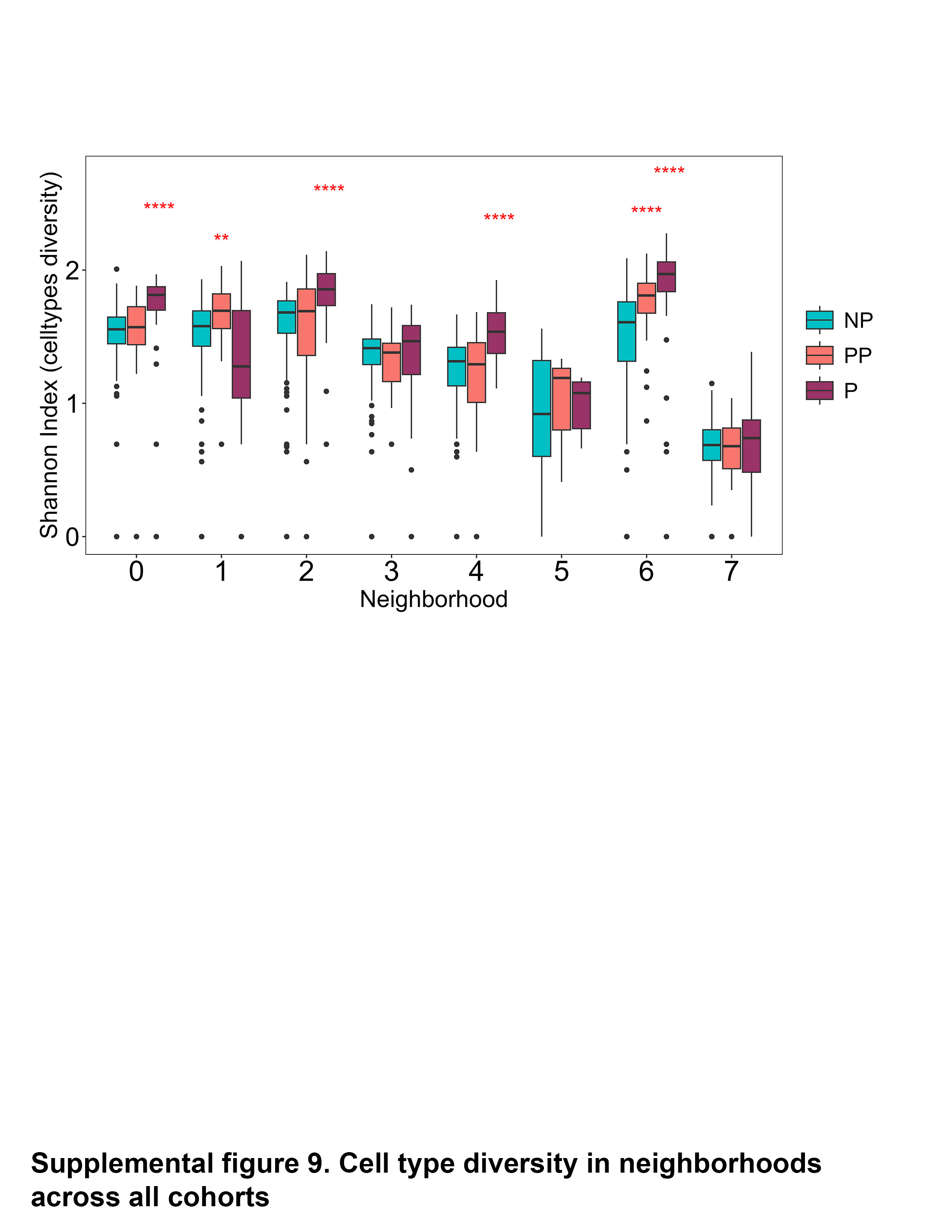

### Supplemental figure 10A

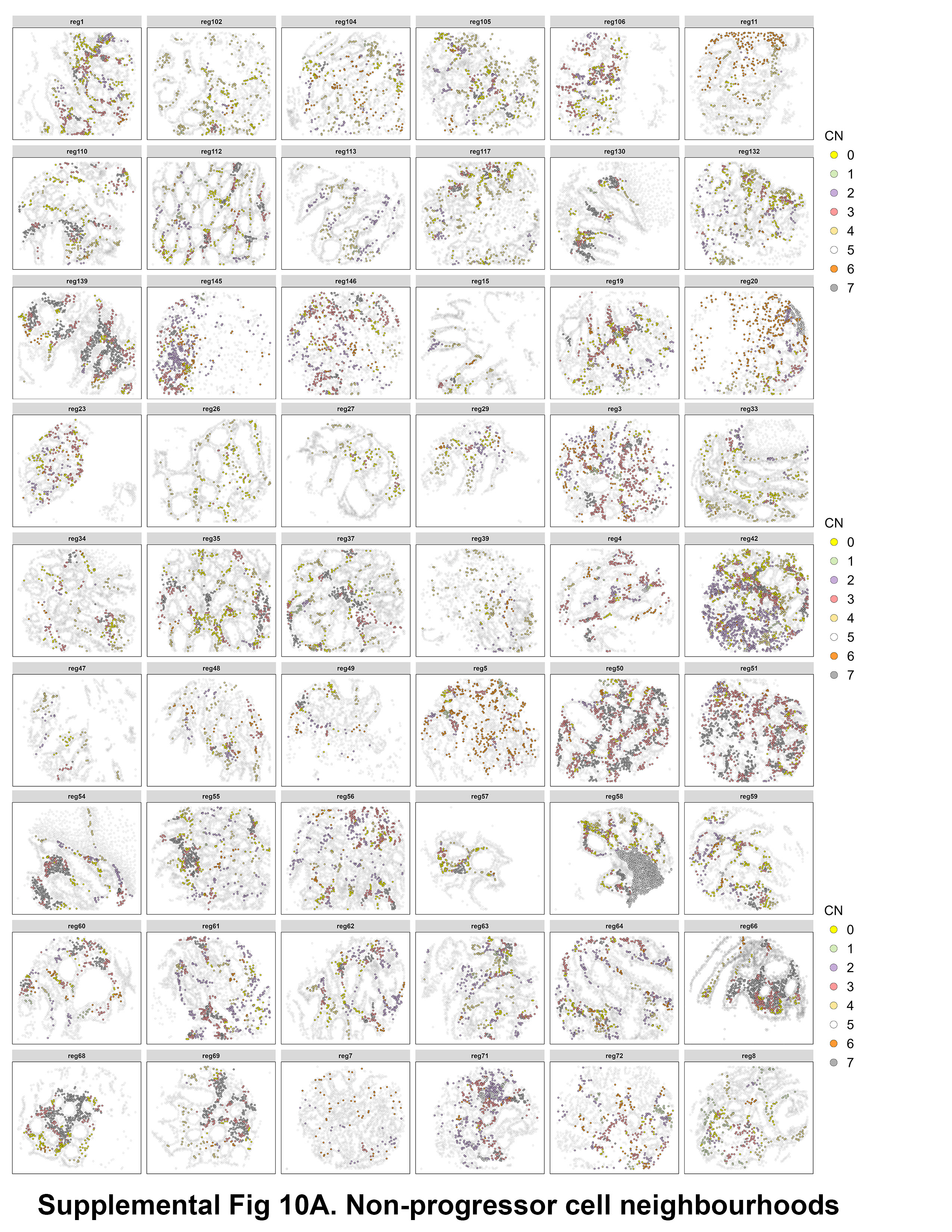

### Supplemental figure 10B

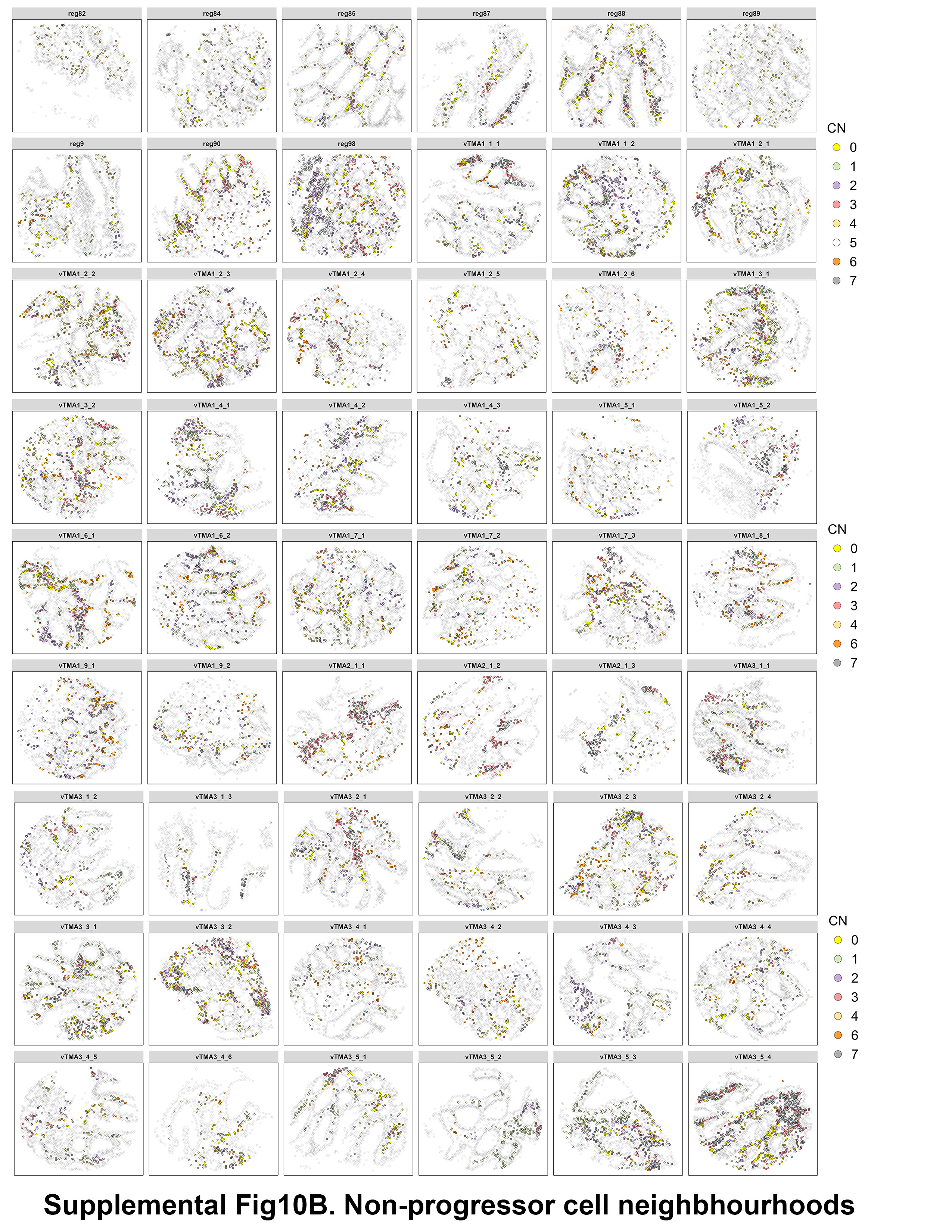

### Supplemental figure 10C

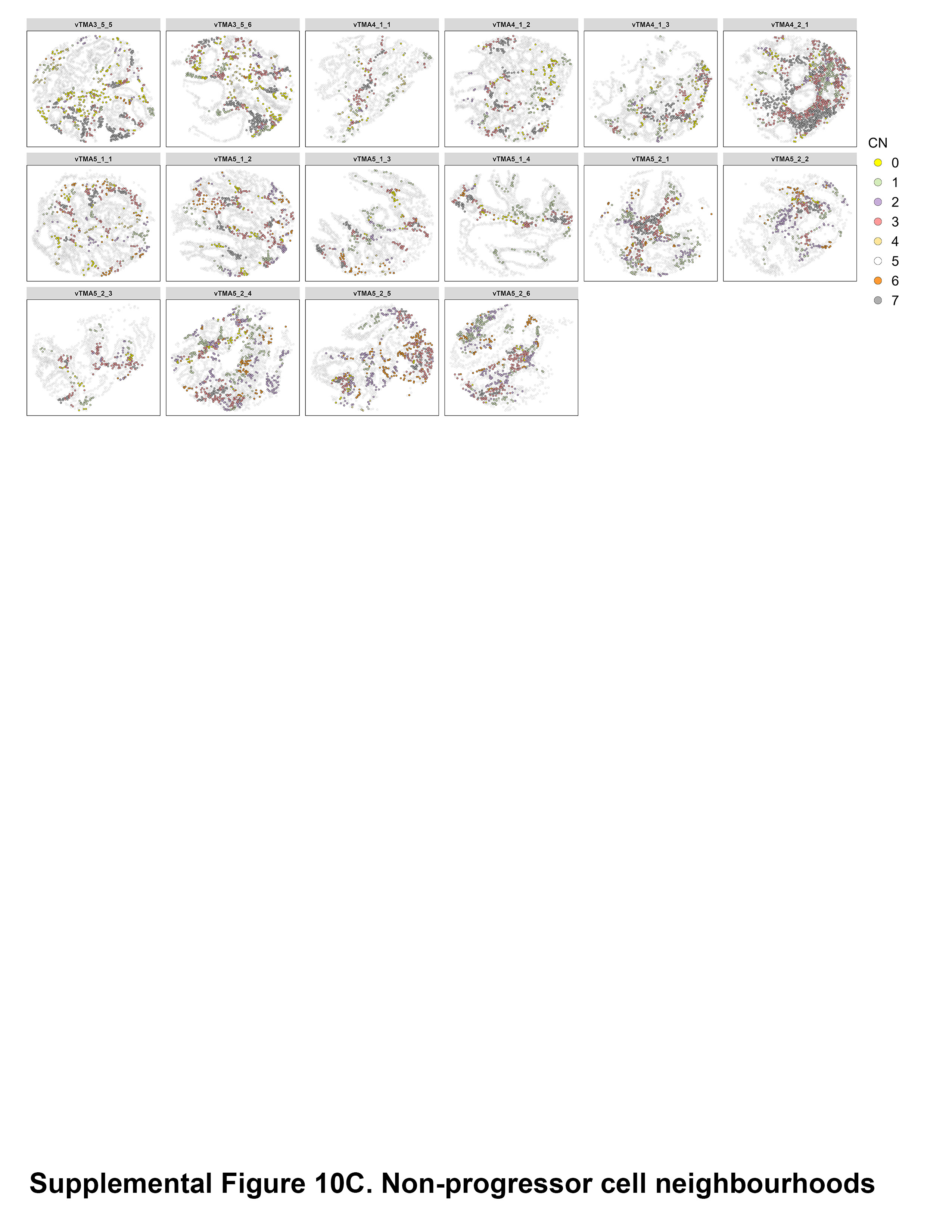

### Supplemental figure 10D

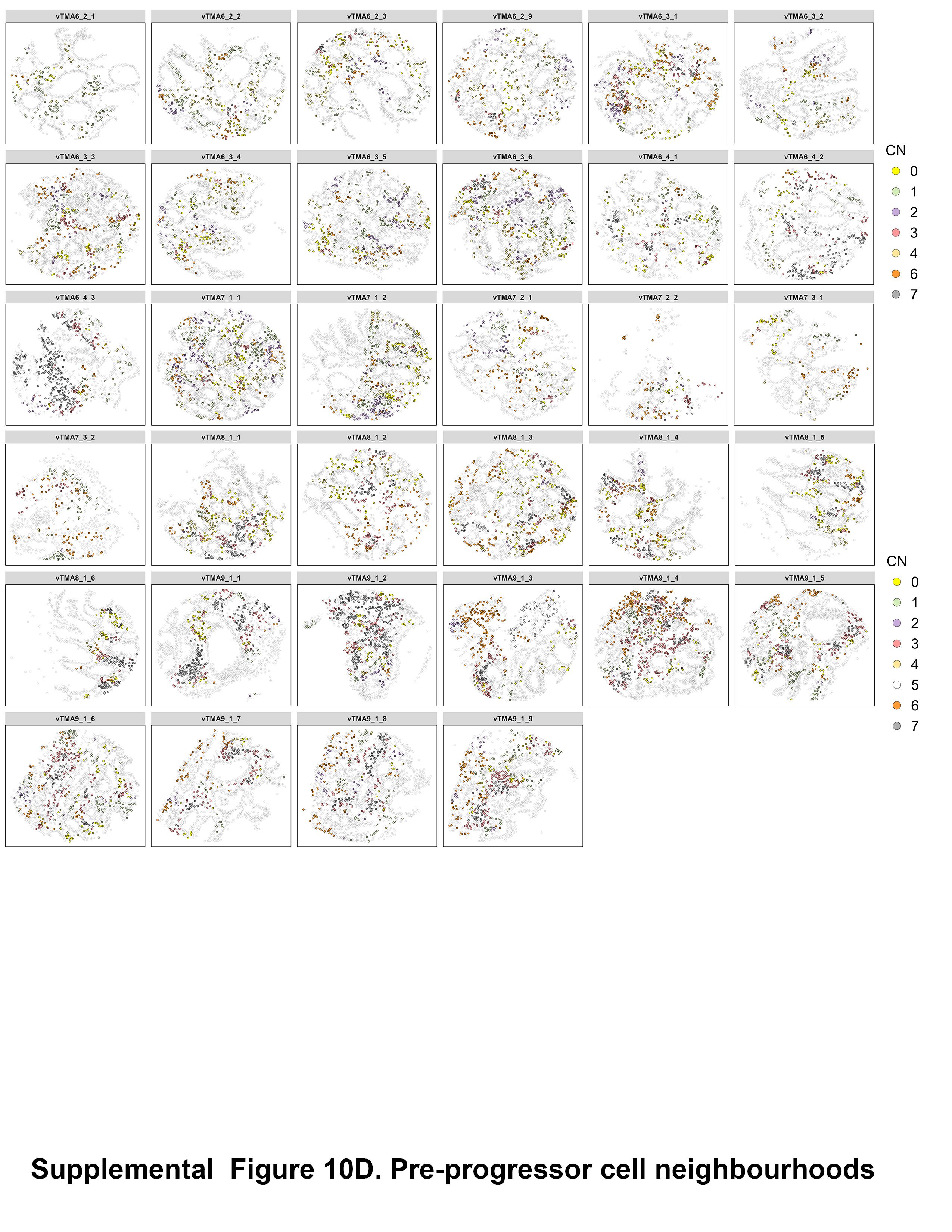

### Supplemental figure 10E

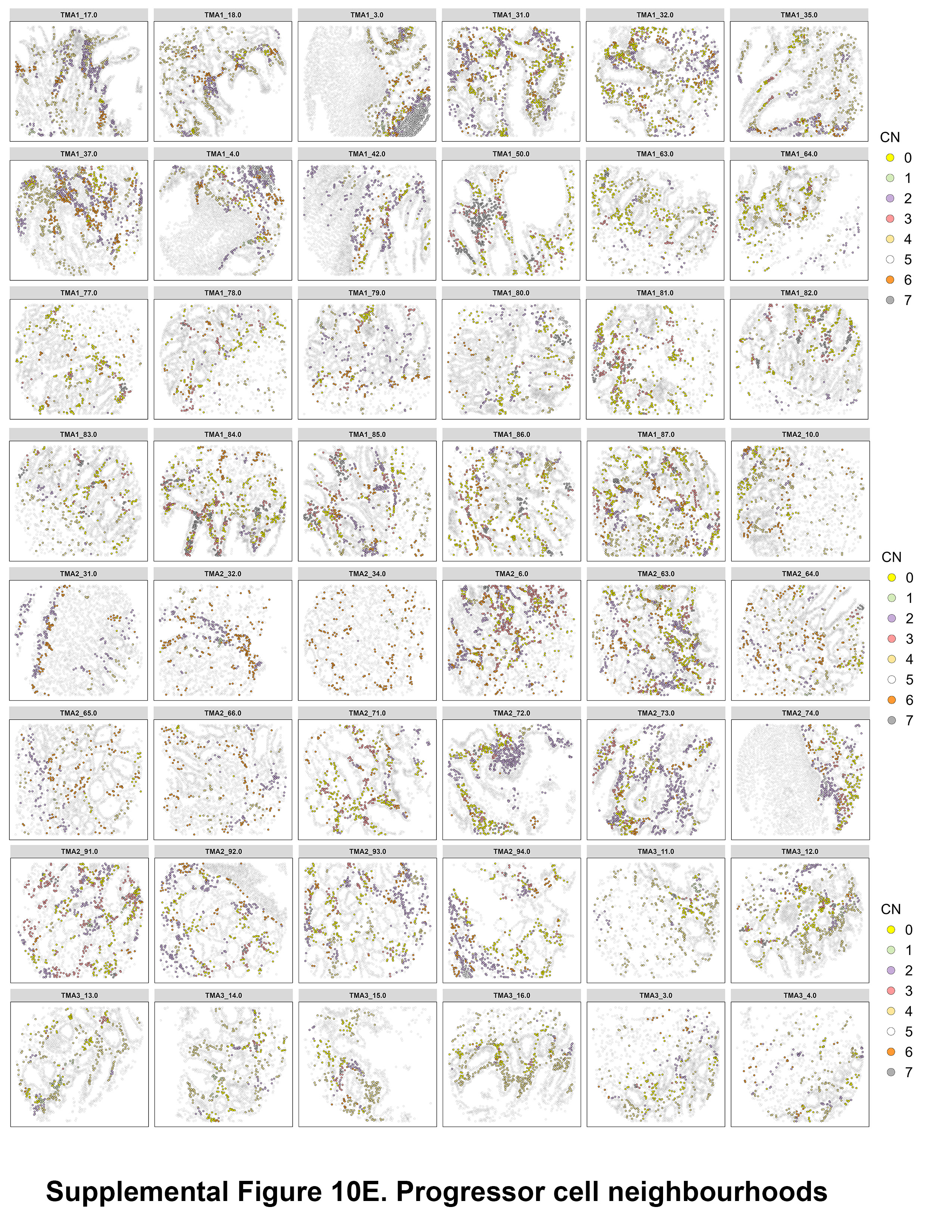

### Supplemental figure 11

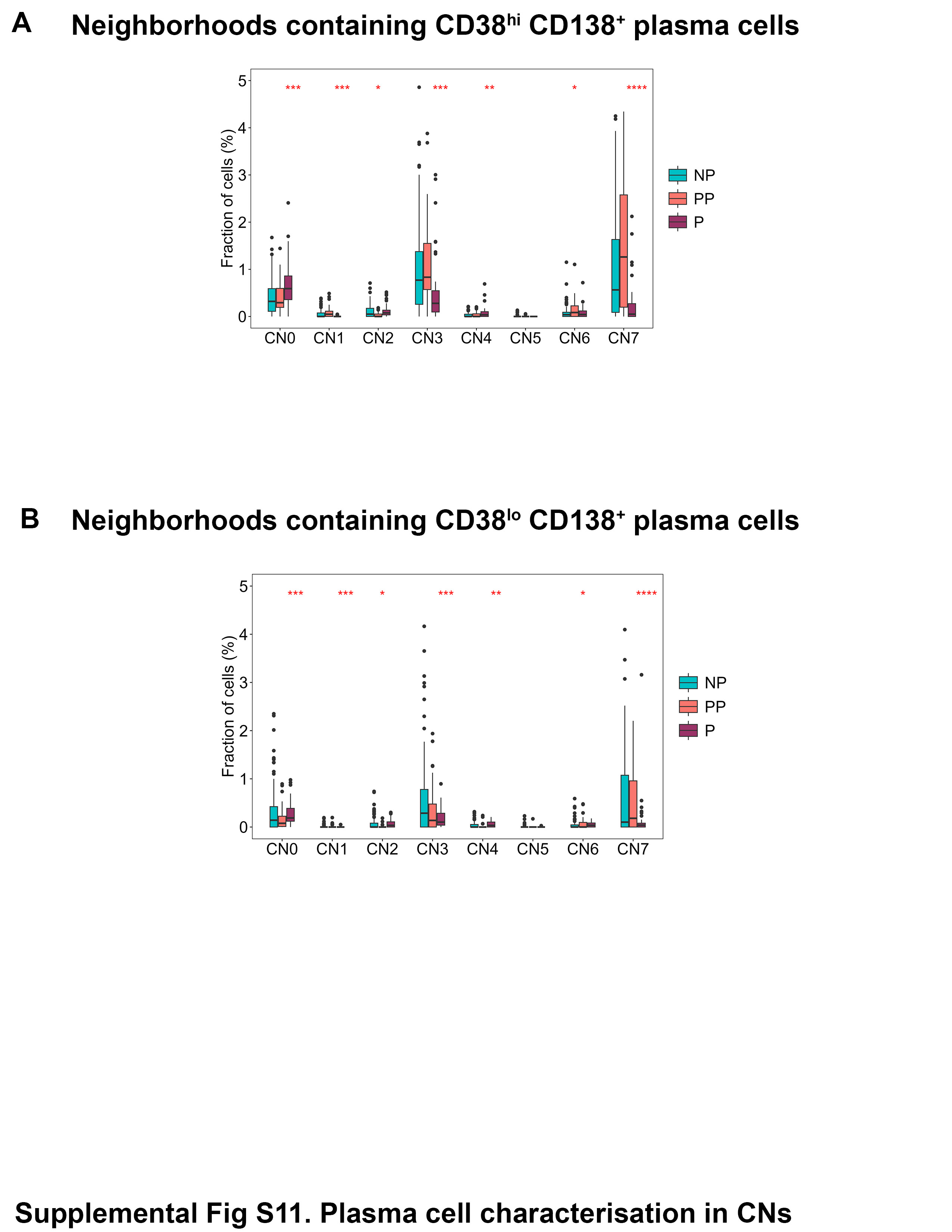

### Supplemental figure 12

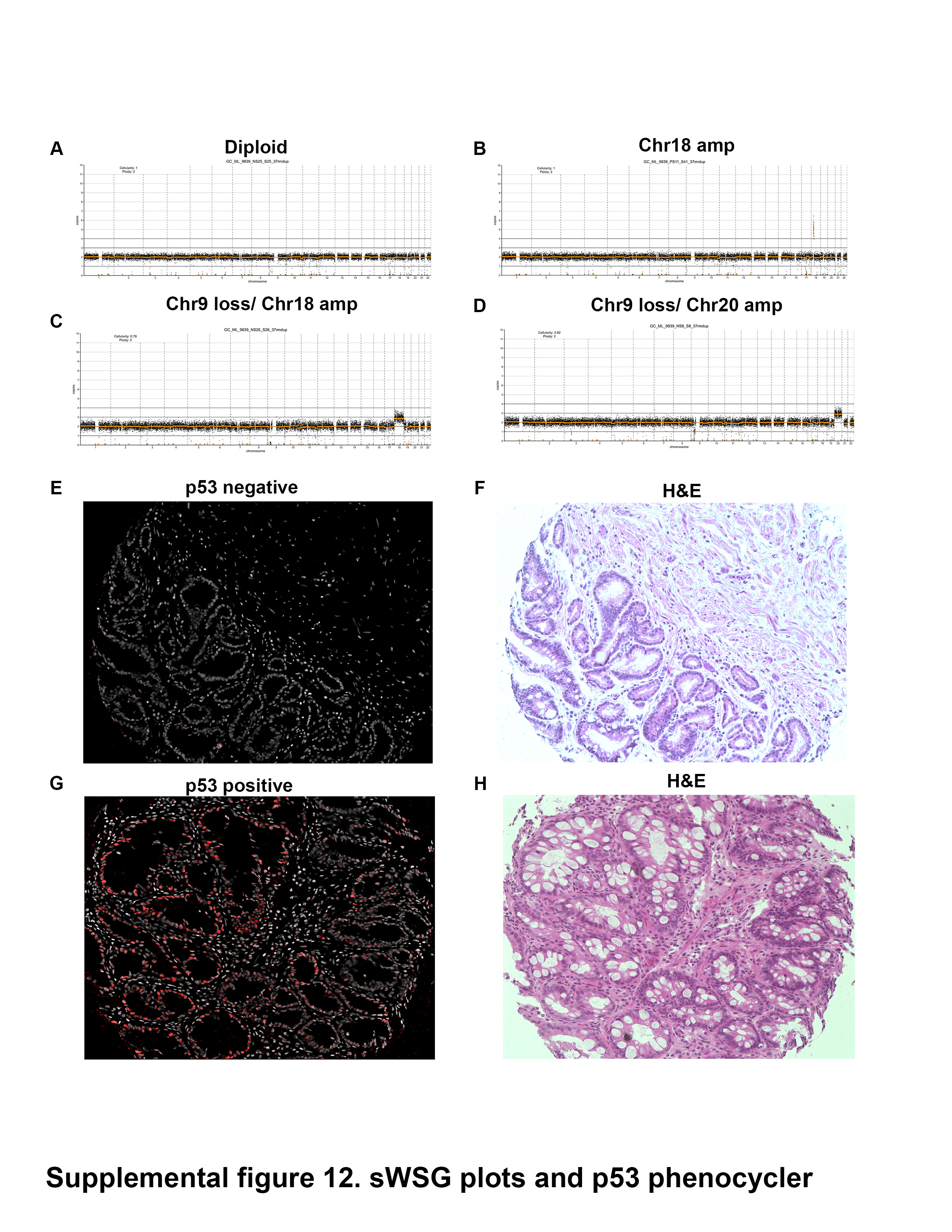

### Supplemental figure 13

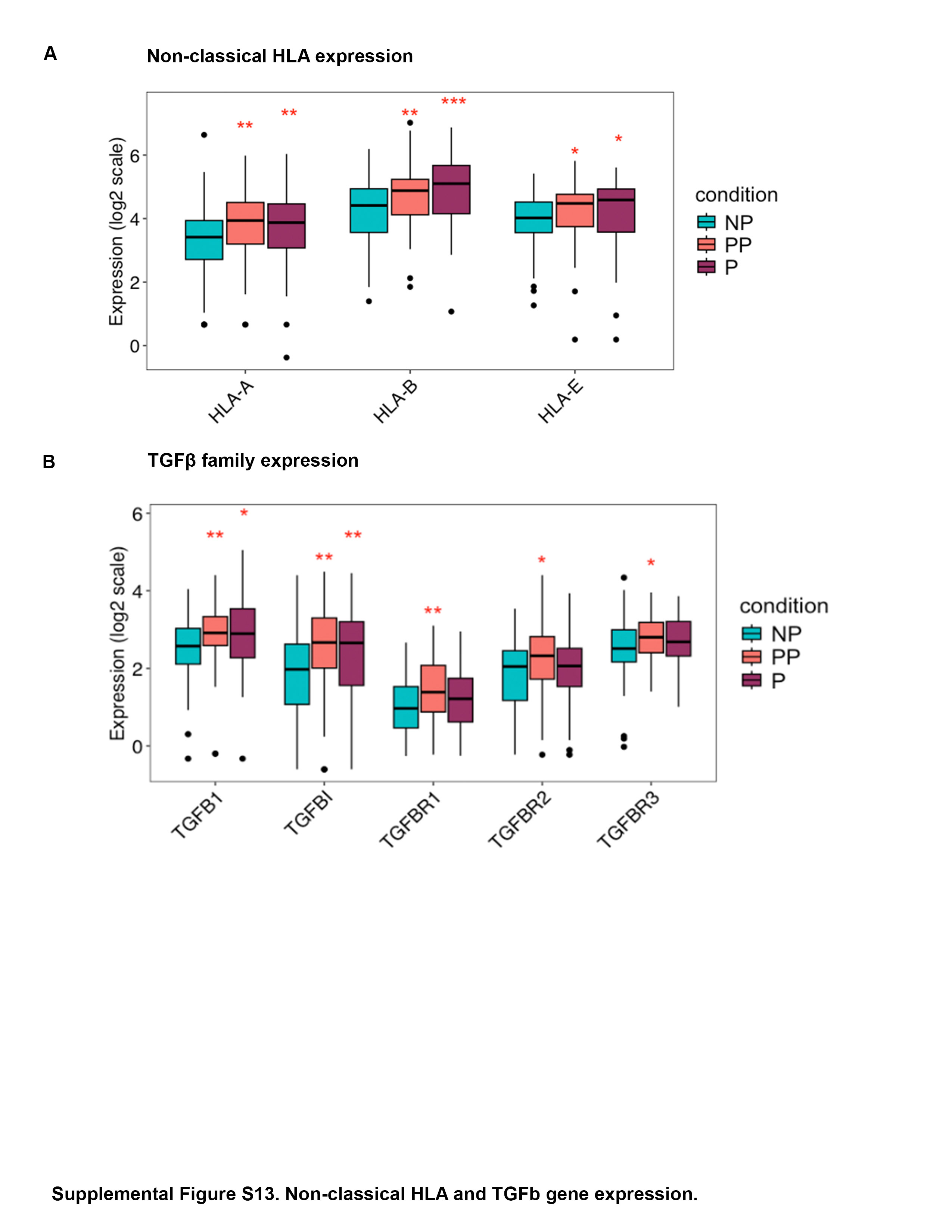
